# Consensus across limbic areas predicts motivational disengagement

**DOI:** 10.64898/2026.07.28.741314

**Authors:** Diana C. Burk, Bruno B. Averbeck

## Abstract

Motivational disengagement is widely attributed to low expected reward, yet animals and humans disengage even when reward expectations are constant. This implies a latent mechanism beyond reward. After accounting for expected reward, we found a neural signal across multiple limbic areas that predicted disengagement. As animals disengaged, neural activity across areas converged onto a shared, low-dimensional axis. Consensus across areas predicted whether disengagement persisted, suggesting why single brain area perturbations have produced inconsistent effects. We showed that engagement and disengagement operated as distinct attractor states whose strength scaled with expected reward, revealing that value modulates, but does not determine, motivational state. These findings reframe motivation as a collective computation and suggest that effective interventions for motivational deficits may require targeting multiple areas simultaneously.

## Introduction

Motivation governs whether goal-directed behavior is initiated and sustained. It is disrupted across psychiatric illnesses, manifesting as anhedonia in depression, avolition in schizophrenia, and engagement lapses in attention-deficit disorders (*1–4*). Although motivation is conserved across species and central to everyday behavior, a unified circuit-level account of how the brain mediates transitions between engagement and disengagement has remained elusive.

When motivation is manipulated through a specific physiological drive or reward, neural signals reflecting motivational state are difficult to dissociate from those encoding the value of the reward at stake (*5, 6*). Furthermore, neural activity is often analyzed during task execution rather than in the periods beforehand that govern task engagement. Consequently, it remains unknown how brain circuits configure motivational state beyond what is predicted by reward, in the moments before behavior is initiated.

The neural substrates of motivational control include a set of interconnected structures — the ventral striatum, ventral pallidum, lateral hypothalamus, amygdala and insula — each with established roles in reward valuation, arousal, interoception (e.g. satiety), and affective processing (*7–17*). These structures are anatomically coupled through dense reciprocal connections and are collectively referred to as limbic motivational circuitry (*12, 18–20*). Each structure has been studied individually for its contribution to motivated behavior. But whether their collective coordination mediates transitions between engagement and disengagement remains unresolved.

Here we recorded simultaneously across five limbic areas while rhesus macaques performed a reinforcement learning task, the structure of which generated fluctuations between engagement and disengagement. We found that disengagement was reflected in a coordinated re-organization of population activity across limbic regions, with each area shifting collectively onto a shared motivational axis. In addition, the degree of consensus, or cross-area commitment to the disengaged state, predicted whether disengagement would persist. Network dynamics analyses revealed that the limbic circuit represents motivational engagement and disengagement as distinct attractor states, the stability of which scaled with expected reward. Our results show that motivational state is carried in the coordinated dynamics of the neural circuit, with implications for understanding why single-target interventions have produced inconsistent clinical effects.

## Results

### A token bandit task leads to fluctuations in motivation

To examine natural transitions between engagement and disengagement, we had two rhesus macaques (Monkey W, n=30 sessions; Monkey G, n=39 sessions) perform a dual-fluid tokens bandit task (Fig 1A, fig. S1). Tokens served as symbolic reinforcers that accumulated visibly across trials and were later exchanged for fluid rewards at cashout, which occurred every four to six trials (*21–23*). On each trial, the monkey earned tokens by choosing between two of the presented images. Trials in which the monkey failed to acquire or hold fixation, or failed to acquire or hold a cue image, were considered aborts, and the trial immediately repeated with no change in token count.

**Fig. 1.**
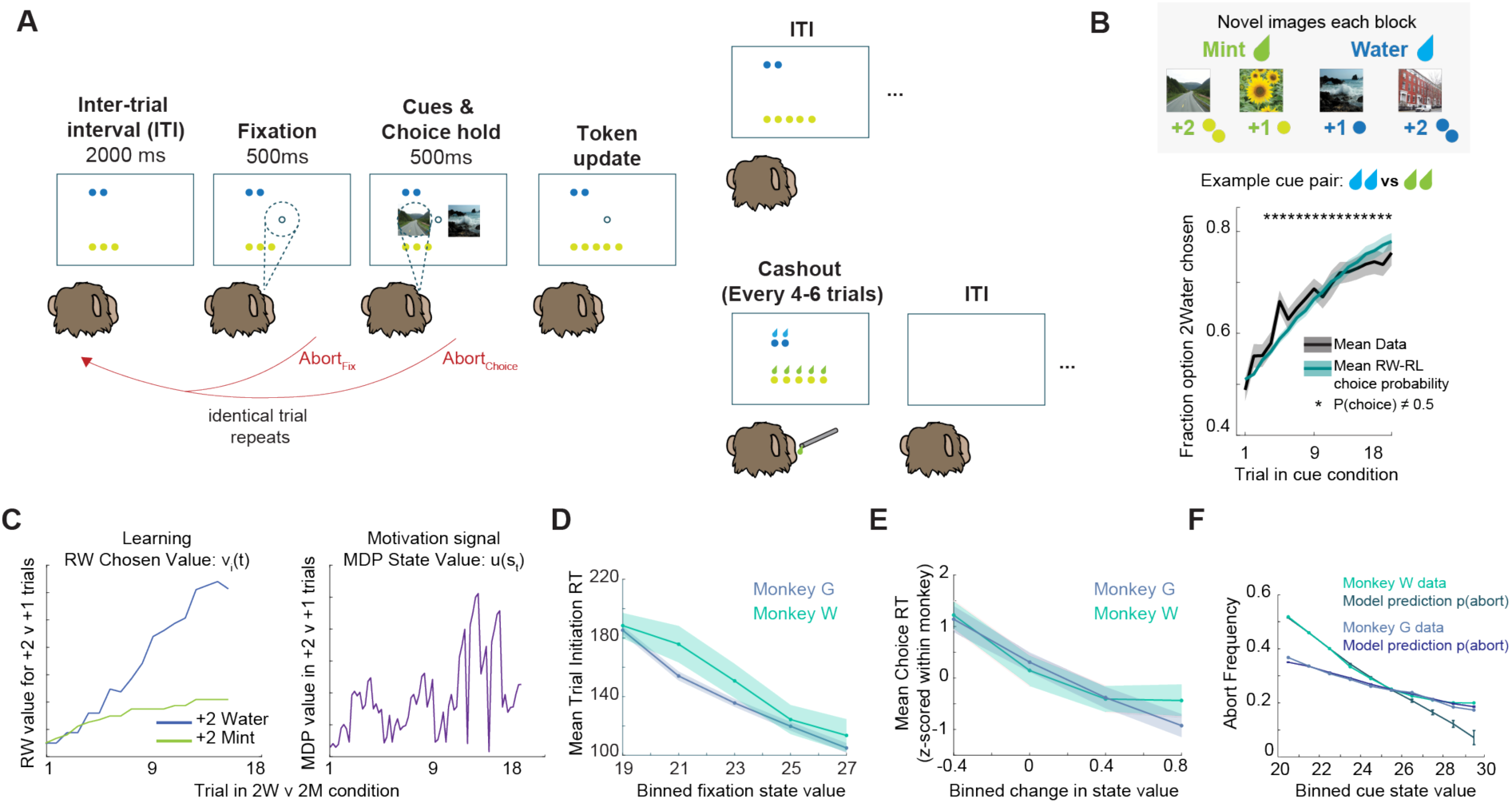
A token-based task reveals behaviorally structured motivational disengagement. **(A)** Task schematic. Two monkeys chose between visual cues associated with mint or water tokens. Successful choices earned tokens on 80% of trials. Tokens accumulated across trials and were exchanged for fluid during cashout, then reset to zero. Failure to acquire or hold fixation or the cue constituted an abort; the trial repeated immediately with no token change. Inter-trial interval (ITI), 2000 ms. **(B)** *Top:* Four novel images were introduced every block of 108 trials and each image was associated with +1 or +2 mint or water tokens. *Bottom:* Example of choice behavior in one of the six task conditions: 2 Water vs. 2 Mint demonstrating learning of cue values. Mean fraction of trials on which the 2 Water option was chosen across trials averaged across n=69 sessions from both monkeys (black), overlaid with the choice probability computed from the Rescorla-Wagner (RW) model fit separately to each session (teal). Asterisks indicate trials at which choice probability differed significantly from chance (BH-FDR corrected, p < 0.05). Shaded error bars s.e.m. **(C)** RW chosen value (left) and Markov decision process (MDP) state value (right) for the example 2 Water vs. 2 Mint condition. **(D to F)** Behavioral correlates of MDP state value. Monkey W, n = 30 sessions; Monkey G, n = 39 sessions. Shaded error bars s.e.m. across sessions. **(D)** Trial initiation reaction time (RT), **(E)** Choice RT, and **(F)** Abort frequency with MDP model-predicted abort probability.

Each block used four novel images associated with specific token outcomes: two images predicted mint-flavored fluid tokens (+2 Mint or +1 Mint) and two predicted water tokens (+2 Water or +1 Water) (Fig. 1B, top). The cue-token associations had to be learned by trial and error. The six possible image combinations were presented pseudorandomly within each block. The two fluid types were chosen to recruit neural structures sensitive to fluid identity and interoceptive reward state. Monkeys learned to select the higher-value option associated with their preferred fluid (Fig 1B, fig. S2). In conditions where neither option was preferred, choice probability remained near chance and reaction times slowed, a pattern consistent with reduced motivational engagement rather than failed learning (fig. S2).

To track the component of motivational state related to reward, we computed a state value signal using a Markov decision process (MDP) model fit to each monkey’s behavior (*21*). The MDP model integrated the reward-predictive features of the task (i.e. mint token count, water token count, time since last cashout, cue-outcome learning, trial epoch) into the *state value*, which quantifies the monkey’s immediate and future expected reward. A high state value characterizes states where the monkey expects a large reward soon and a low state value characterizes states where expected rewards are smaller or more distant in time. Cue value learning was captured using a Rescorla-Wagner (RW) model. RW chosen value estimates served as one input to the MDP, allowing state value to incorporate trial-by-trial learning alongside token accumulation and cashout timing at sub-trial resolution (Fig 1C).

State value predicted three behavioral signatures of motivation, replicating our prior findings that state value is a behaviorally grounded metric of motivational state (*21*). Monkeys initiated trials faster when expected reward was high, as measured by fixation state value (Monkey W: t(29) = -25.95, p < 0.0001; Monkey G: t(38) = -28.09, p < 0.0001; Fig 1D). Monkeys made faster choices when state value increased more from fixation to cue presentation (Monkey W: t(29) = -7.11, p <0.0001; Monkey G: t(38) = -12.65, p < 0.0001; Fig. 1E). Monkeys aborted trials more frequently when state value was low (Monkey W: t(29) = -8.99, p < 0.0001; Monkey G: t(38) = -6.87, p < 0.0001; Fig. 1F). However, abort probability only captures the average tendency to abort individual trials, not how disengagement unfolds across sequences of abort trials. Because state value remained constant across consecutive aborts, the sequential structure of disengagement was not accounted for by reward expectation alone. This led us to characterize the behavioral and neural transitions from engagement to disengagement.

### Motivational disengagement unfolds as recurring sequences of abort trials

We examined disengagement over sequences of abort trials as an innate behavior related to motivation. Although Monkey W and Monkey G differed in average abort rates across the session (Fig. 2A), both monkeys aborted trials in recurring sequences that were distributed throughout the session (Fig. 2B, 2C) and were minimally affected by cue condition (fig. S3). When we examined the epoch within the trial at which the abort occurred, we found that single aborts (Completed-Abort-Completed Sequence: C-A_1_-C) were concentrated during fixation and choice hold epochs, whereas sequential aborts (Abort-Abort-Abort Sequence: A_1_-A_2_ -…-A_N_) were mostly a failure to acquire initial fixation. This suggests a higher proportion of motor errors in single aborts (fig. S3). Disengagement, which we define as the transition into a sequence of aborts, was therefore a recurring phenomenon, distinct from session-edge effects such as warm-up or fatigue and not accounted for by motor errors alone.

We next asked whether state value at the first abort (C-A_1_) distinguished a transient lapse from a disengagement. We compared the mean inter-trial interval (ITI) state value between trial sequences in which the first abort (A_1_) was followed by a completed trial (recovery: A_1_-C) and those in which the first abort was followed by at least three consecutive aborts (disengagement: A_1_-A-A-A). State value at the first abort was significantly lower preceding sustained disengagement than in single aborts (Fig. 2D). Thus, when expected rewards were higher, the monkeys were less likely to disengage. Because state value remained constant across the abort sequence, however, the depth and duration of disengagement reflected something beyond reward expectation.

Following disengagement, monkeys re-engaged gradually across completed trials (Fig. 2E, F). Trial initiation times decreased significantly from the first completed trial (C_1_) to the second (C_2_) and from the second to the third completed trial (C_2_ to C_3_) in both monkeys, with no further significant change beyond the third completed trial (Fig. 2E). Choice reaction times recovered faster. The significant drop from the first to the second completed trial was not followed by any further significant change (Fig. 2F). Having characterized the behavioral structure of disengagement and its modulation by state value, we then asked how transitions in motivation are represented across limbic areas.

**Fig. 2:**
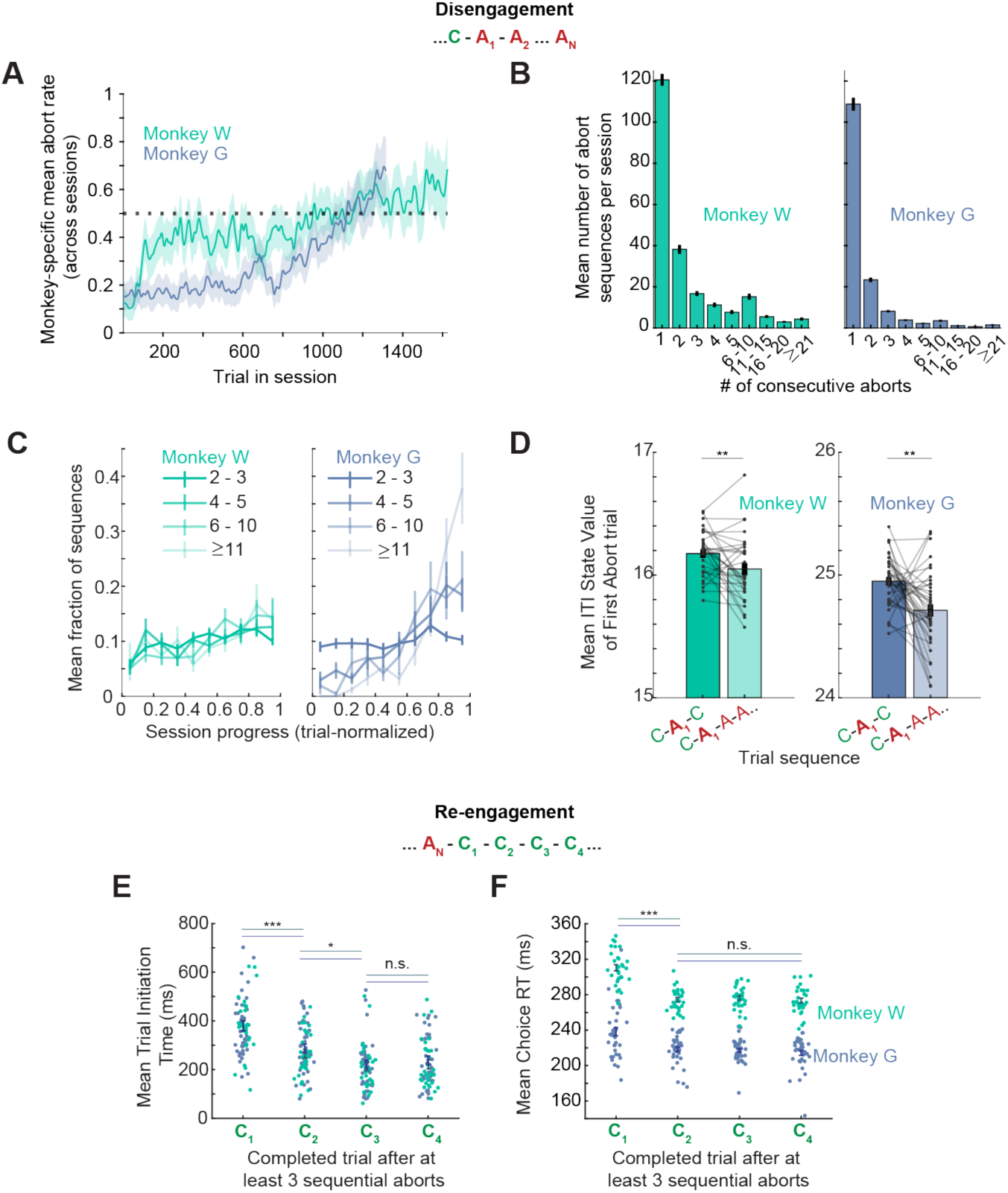
Behavioral signatures of disengagement and re-engagement. **(A)** Mean abort rate across session trials for each monkey. Monkey W shows a gradual increase; Monkey G shows a U-shaped profile. Dashed line, 0.5. **(B)** Mean number of abort sequences per session as a function of consecutive abort count. Single aborts are most frequent; longer sequences occur in both animals. **(C)** Mean fraction of sequences of each length as a function of normalized session progress. **(D)** Mean ITI state value on the first abort of a sequence, separated by subsequent trial engagement (C-A-C or C-A-AA…). (Monkey W: t(29)= -2.94, p=0.006; Monkey G: t(38)=-4.06, p=0.002) **(E and F)** Re-engagement dynamics in behavior after abort runs of at least three consecutive aborts. **(E)** Mean trial initiation time (C1◊C2 Monkey W: t(29) = -8.90, p <0.0001; Monkey G: t(38) = -4.56, p = 0.0002; C2◊C3 Monkey W: t(29) = -3.02, p = 0.011; Monkey G: t(38) = -4.01, p=0.0005; C3◊C4: Monkey W: p = 0.583; Monkey G: p = 0.377). **(F)** Mean choice RT (C1 ◊ C2 :Monkey W: t(29) = 8.49, p < 0.0001; Monkey G: t(38) = 4.37, p = 0.0002; C2 ◊ C3 : Monkey W: p = 0.86; Monkey G: p = 0.691. Individual session means shown as dots; shading and error bars, s.e.m. across sessions.

### Limbic areas encode motivation prospectively during the inter-trial interval

To examine the neural representation of motivational state, we performed simultaneous multi-site recordings spanning five interconnected structures: the ventral striatum (VS), ventral pallidum (VP), lateral hypothalamus (LH), amygdala (Amyg), and anterior insula (Ins) (Fig. 3A, B). We used a previously published targeting approach with semi-chronic guide tubes and multi-contact linear arrays (Fig. 3C, fig. S4, (*23*), which yielded 7,759 total units (VS: 1,236; VP: 1,756; LH: 1,337; Amyg: 2,006; Ins: 1,424; table S1). Across areas, units were modulated by all task features, including state value, token outcomes, cue identity, and cue value throughout task epochs (fig. S5, fig. S6).

**Fig. 3:**
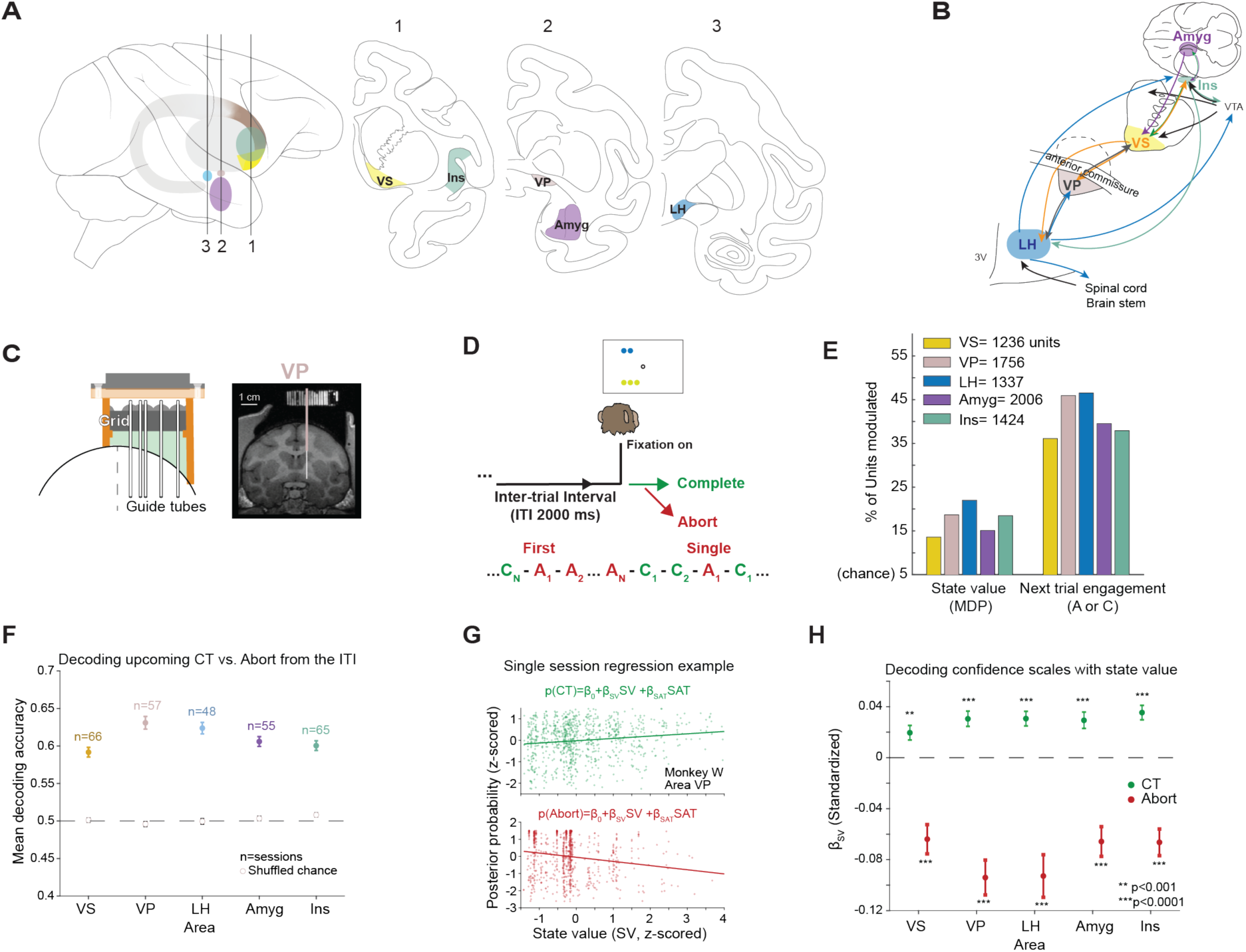
Neural activity across limbic areas predicts motivational state. **(A)** Recording locations for five areas: ventral striatum (VS), ventral pallidum (VP), lateral hypothalamus (LH), amygdala (Amyg), and anterior insula (Ins). **(B)** Schematic of anatomical connectivity among recorded structures. **(C)** MRI-guided semi-chronic recording approach illustrated for VP (Monkey W); electrode trajectory shown as colored overlay. Representative recording locations for all areas in fig. S4. **(D)** Schematic of ITI analysis window and abort sequence notation describing first and single aborts. **(E)** Percentage of units responsive to State value or Next trial engagement (A or C), during the ITI. Chance level 5%. **(F)** Cross-validated decoding accuracy for upcoming abort versus completed trial from mean ITI population activity, by recording area, sessions pooled across monkeys. n indicates number of sessions. Open circles: shuffled-label chance; dashed line: theoretical chance at 0.5. **(G)** Single-session example (Monkey W, area VP) showing posterior probabilities regressed on state value (SV) and a time-in-session or satiety proxy (SAT), separately for CT and abort trials. **(H)** Standardized SV regression coefficients (βSV) across areas for CT and abort trials; Full statistics in table S2. Error bars: s.e.m. across sessions.

Because state value was encoded throughout the trial, including during the ITI, we asked whether population activity during the ITI would predict behavioral engagement before the fixation spot appeared (Fig. 3D). To characterize ITI encoding, we used a separate analysis to quantify the fraction of units significantly modulated by features available at that epoch, including outcome-related information from the preceding trial, state value, and upcoming trial engagement (C or A) (Fig. 3E, fig. S7, fig. S8). Across areas, 10-25% of units were modulated by state value during the ITI and a substantially larger proportion of units (35-50% per area) were modulated by upcoming trial engagement (Fig. 3E, fig. S8A). The two populations in each area were largely independent, with the extent of shared coding exceeding chance by only 15-19% in VP and Amyg, indicating that state value and engagement predictive signals were mainly carried by distinct neuronal populations (fig. S8B). This demonstrated that there were signals at the population level predicting engagement that extended beyond state value.

Although the single-unit analyses revealed that individual neurons carried engagement-predictive information, they did not address whether the information was sufficient to predict behavior. Using linear SVM classifiers trained on mean ITI population activity of all units for each recording session, we decoded whether the upcoming trial was completed or aborted. Decoding accuracy exceeded chance in all five areas (Fig. 3F), indicating that activity during the ITI was sufficient to predict the monkey’s task engagement before behavioral onset. We also characterized the time-course of this signal across the 2000 ms ITI window to examine the onset of a disengagement signal, but performance was flat across bins in all areas, suggesting that the signal reflected a sustained population state without abrupt onset during the ITI (fig. S9). Thus, each area carried sufficient information during the ITI to predict the monkey’s upcoming behavioral engagement.

We investigated how motivational state evolved across sequences of trials by converting trial-by-trial SVM classification scores into posterior probabilities, which provided a graded measure of the classifier’s confidence in each outcome (C or A) for each ITI. Each trial was assigned the posterior corresponding to its true outcome: p(abort|PR) for abort trials and p(completed|PR) for completed trials (where PR indicates population response). Values near 1, which reflect neural activity far from the classifier’s decision boundary, indicated high confidence in the predicted class. Values near 0.5, which reflect neural activity on the decision boundary, indicated chance-level discrimination. Across all five areas, p(abort|PR) increased and p(completed|PR) decreased with decreasing state value, indicating that posteriors extracted from the decoder tracked reward-related motivational state (Fig 3G, H). We also ruled out time-in-session as an alternative explanation for the posterior signal. A time-in-session predictor, included to capture residual temporal structure across the session, had inconsistent and weak relationships with posterior probability (fig. S10, table S2), confirming that the motivation signal captured by the decoder could not be accounted for by time-in-session. These posteriors provided a trial-level measure of motivational state that we used to track how areas transitioned between engagement and disengagement.

### Limbic areas track motivational state asymmetrically across disengagement and re-engagement

We had already observed that when monkeys disengaged, aborts shifted to earlier trial epochs. In abort sequences monkeys mostly did not initiate trials. During re-engagement, trial initiation and choice times recovered gradually across completed trials, suggesting a gradual process underlying both transitions. We next asked whether neural circuit dynamics during disengagement and re-engagement were abrupt or gradual. To examine these possibilities, we computed the slopes of the posterior probability across trial positions and measured whether the posteriors increased or decreased over trials. These slopes revealed an asymmetry between disengagement and re-engagement.

Leading into disengagement, the population state in the completed trial run preceding the first abort showed no significant slope in any area (Fig. 4A; table S3), indicating that the neural activity was not building towards the first abort across completed trials. However, once the abort sequence began, the posteriors rose across abort trials in all five areas (Fig. 4A, left; table S3). In contrast, during re-engagement, posteriors across all areas declined significantly across the late abort run, reflecting a shift toward the engaged state that preceded behavioral re-engagement. The slope across the first three completed trials after an abort sequence was also significant in all areas, showing gradual re-engagement on both sides of the transition.

We further asked whether the population decoding reflected a uniform shift in mean rates or a heterogeneous signal in which individual neurons changed in consistent but opposing directions across the transition. Mean firing rates of engagement-modulated units in VP and LH shifted significantly at transition boundaries, with VP leading during both disengagement onset and re-engagement (fig. S11), consistent with a prominent role for VP across both transitions. These mean rate shifts were restricted to VP and LH, however, suggesting that population-level decoding captured motivational state changes that were not reflected in mean firing rates across all areas.

### Cross-area consensus at the onset of disengagement predicts its persistence

The progression of disengagement was reflected in the change in posteriors over the first few abort trials. But whether the neural state preceding disengagement carried information about what followed could not be resolved by averaging across sequences. We examined the population state at the first abort directly and asked whether its position in a low-dimensional state-space predicted the duration of disengagement that followed. We characterized the geometry of ITI activity preceding first and single aborts (A_1_), later aborts (A_2+_), and completed trials (C) using the top three principal components estimated on the ITI population activity. Consistent with our decoding results, average activity in completed trials and later aborts occupied non-overlapping regions of this low-dimensional subspace, with PC1 providing the primary axis separating the two trial types (Fig. 4B). First aborts and single aborts occupied an intermediate position between the completed trial and later abort clusters. This raised the question of whether the degree of the displacement in the first abort trial predicted whether disengagement would persist.

If the degree of displacement at the first abort predicted the depth of disengagement that followed, then first abort trials farther from the engaged population state would be expected to initiate longer abort runs (Fig. 4C, left schematic). To test this, we computed the distance of each first abort trial’s population activity from the completed-trial session-level mean across the first three PCs, then categorized first abort trials by the length of the abort sequence they initiated (A_1_C_1_ through A_1_AAAA+). Mean distance increased with future abort sequence length across all five areas (Fig. 4C, right), with the strongest effects in VP and LH. The consistent displacement effect across all five areas raised the possibility that what mattered for predicting disengagement was not the magnitude of displacement in any individual area, but the number of areas that were jointly displaced beyond the engaged state.

Under this area consensus hypothesis, the number of areas displaced beyond the engaged state at the first abort (*vote fraction*) would predict whether disengagement persisted. First abort trials in which more areas were displaced from the engaged state (e.g. 4/5 or 5/5 areas) would be expected to be followed by another abort (A_2_), while trials in which only one or two areas were displaced (e.g. 1/5 or 2/5 areas) would be expected to be followed by a completed trial (C_1_) (Fig. 4D, left). To test this hypothesis, we converted the trial-level A_1_ distance into a binary vote per area: 1 if the distance exceeded a median threshold and 0 otherwise. Summing votes across simultaneously recorded areas gave a vote fraction for each A_1_ trial. Across sessions, p(next trial=A_2_) rose with vote fraction and shuffling the votes within each area abolished this effect (Fig. 4D, right). The effect also held in recording sessions with three or four areas (fig. S12A) and the joint vote distributions departed from the binomial null expected if votes were independent across areas (fig. S12B). Shuffling next-trial outcomes within sessions also abolished the effect (null mean β_VF_ = 0.005, SD 0.07, p<0.001). Disengagement was therefore predicted by coordinated displacement across the network, rather than displacement in a single area.

**Fig. 4.**
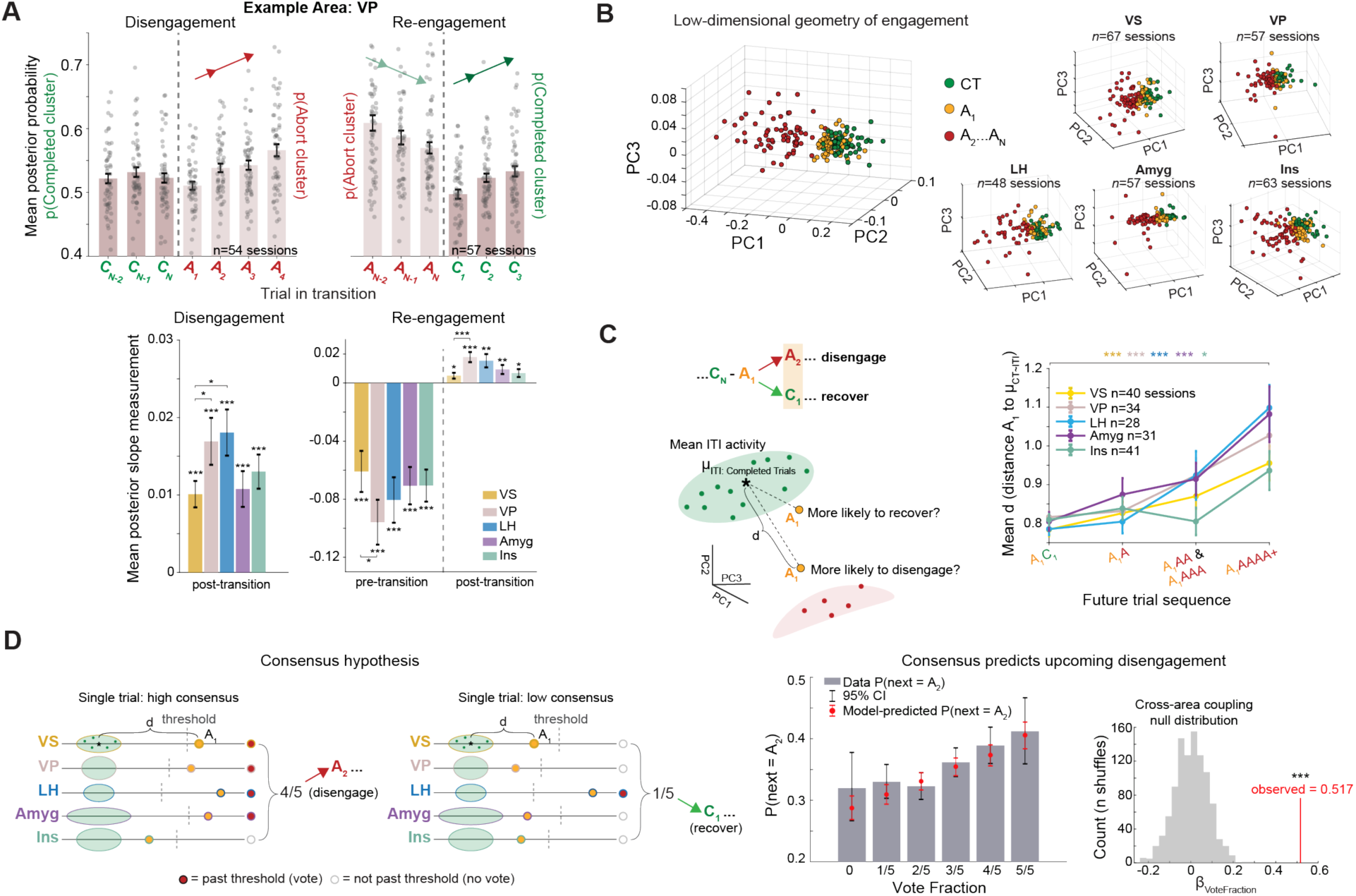
Population geometry and cross-area consensus predict disengagement persistence. Data pooled across monkeys. **(A)** *Top:* Mean posterior probability across sequential trial positions during disengagement (left) and re-engagement (right), shown for VP. Dots: individual session means; error bars: s.e.m. across sessions. *Bottom:* Mean posterior slope measurements across areas. Left: abort-run slopes; VP and LH largest (VP>VS: t(115)=2.030, p=0.022; LH>VS: t(107)=2.455, p=0.016). Right: pre-CT slopes (all areas p<0.0001) and CT-run slopes (VS: t(65)=2.500, p=0.015; VP: t(56)=5.096, p<0.0001; LH: t(47)=3.330, p=0.003; Amyg: t(54)=3.011, p=0.006; Ins: t(64)=2.508, p=0.015); VP steepest recovery (VP<VS (abort): t(121)=1.671, p=0.049; VP>VS (completed): t(121)=3.269, p=0.0007). Full statistics in table S3. **(B)** Session-mean ITI population activity projected onto the first three principal components (PC1, PC2, PC3), by trial type: completed trials (green), first and single aborts (yellow), later aborts A2-AN (red). Each point: session mean. **(C)** *Left:* Schematic of A1 distance metric (d) to session engaged cluster centroid (μITI:CompletedTrials) across the first three PCs, grouped by future sequence length. *Right:* Mean A1 distance by future sequence length; shortest vs. longest category: VS: t(39)=4.016, p=0.0004; VP: t(33)=4.623, p=0.0001; LH: t(27)=4.845, p=0.0001; Amyg: t(30)=3.723, p=0.001; Ins: t(40)=2.523, p=0.016. Error bars: s.e.m. **(D)** *Left:* Consensus hypothesis schematic; high-consensus (4/5 areas displaced ◊ A2) and low-consensus (1/5 areas displaced ◊C1). *Middle:* p(next=A2) as a function of vote fraction, overlaid with GLMM predictions and 95% CI; βVoteFraction=0.517, SE=0.07, p<0.0001; n=6180 trials with all five areas recorded. *Right:* Null distribution of βVoteFraction from within-area shuffles of vote fractions; mean βVoteFractionNull=0.001, p<0.001.

### Disengagement shifts limbic populations onto a shared low-dimensional axis

Prior analyses established that each area tracked motivational state but left open whether areas transitioned independently or as a coupled system. To quantify how inter-area coordination changed across the five areas during disengagement and re-engagement transitions, we used the posterior probabilities from decoding, which enabled us to examine inter-area correlations beyond pairwise interactions. We computed the correlation matrix of the per-area posterior probabilities for each trial in the disengagement and re-engagement transitions (fig. S13 A,B). This characterized the coupled population dynamics representing motivation across the five areas. We summarized the structure of the correlation matrix using the determinant (specifically 1-det(R)) of the matrix, which increases as inter-area correlations increase. A higher value of 1-det(R) indicated that the areas were more coherent.

Inter-area correlation was stable across the last completed trial before disengagement onset (C_N_◊A_1_), then increased significantly from the first abort to the second (A_1_ → A_2_) and continued to increase through the third abort (A_3_)(Fig. 5A, left). During re-engagement, inter-area correlation dropped abruptly from the last abort to the first completed trial (A_N_ ◊ C_1_) but did not change further across subsequent completed trials (Fig. 5A, right). The stability of inter-area correlation into the first abort preceding disengagement was consistent with the intermediate positions of first aborts (A_1_) in the population geometry (Fig. 4B) and suggests that network-level commitment to disengagement crystallized only after the first abort.

Among pairwise area correlations, every pair that increased significantly during disengagement involved VP, with the single exception of LH-Ins (Fig. 5B). This indicated that the progressive inter-area coordination was disproportionately organized around VP. Individual posterior variance increased in parallel across the disengagement sequence (fig. S13C), confirming that rising inter-area correlations reflected genuine co-variation rather than compression of the dynamic range or a floor effect.

Increasing pairwise correlations between areas could reflect coupling among many independent dimensions or a shift onto a single shared axis. To distinguish between these possibilities, we examined the dimensionality of the coordination. PCA on the five-area posterior correlation matrix revealed that the variance explained by PC1 (PC1_PCM_, PCM=Posterior Correlation Matrix) increased progressively during disengagement (Fig. 5C), indicating that the multi-area signal became increasingly one-dimensional. Variance explained by PC1_PCM_ dropped sharply at re-engagement onset (A_N_ ◊ C_1_, Fig. 5C, right) and remained low across subsequent completed trials, indicating that the inter-area correlation structure reset abruptly and did not re-emerge gradually. This was distinct from the gradual recovery of within-area posterior values and reaction times across completed trials (C_1_ through C_3_, Fig. 4A, Fig. 2E, F), suggesting that the network-level effect preceded individual area effects and behavior. Across both engaged and disengaged states, all five areas loaded positively and with similar magnitude onto this shared PC1_PCM_ axis (fig. S13D), indicating that disengagement did not fractionate the circuit into opposing subgroups but instead entrained areas together along a common motivational axis.

### State value modulates the depth of attractor basins along the shared motivational axis

Because areas moved together along a shared motivational axis, we asked whether population dynamics along this axis were governed by attractor-like fixed points or if activity drifted around each motivational state’s mean activity pattern. Under the attractor hypothesis, each motivational state would have a location in state space toward which neural activity is continuously pulled (Fig. 5D, left). To test for the existence of fixed points in the neural data, we fit a linear dynamical systems model (*24*) to the pooled five-area PC1 projections separately for completed trials (C) and late abort trials (A_2+_) on a per-session basis (Fig. 5D, left). For each group of trials, the model estimated a fixed point (x_0_), the value toward which population activity was attracted when undriven, and a retraction coefficient (α), which quantified the rate at which activity was pulled toward the fixed point. Higher α values indicated stronger attraction toward the fixed point and thus a deeper basin. The two parameters capture separate properties of a dynamical state, its mean activity pattern and the strength with which that pattern is maintained.

**Fig. 5:**
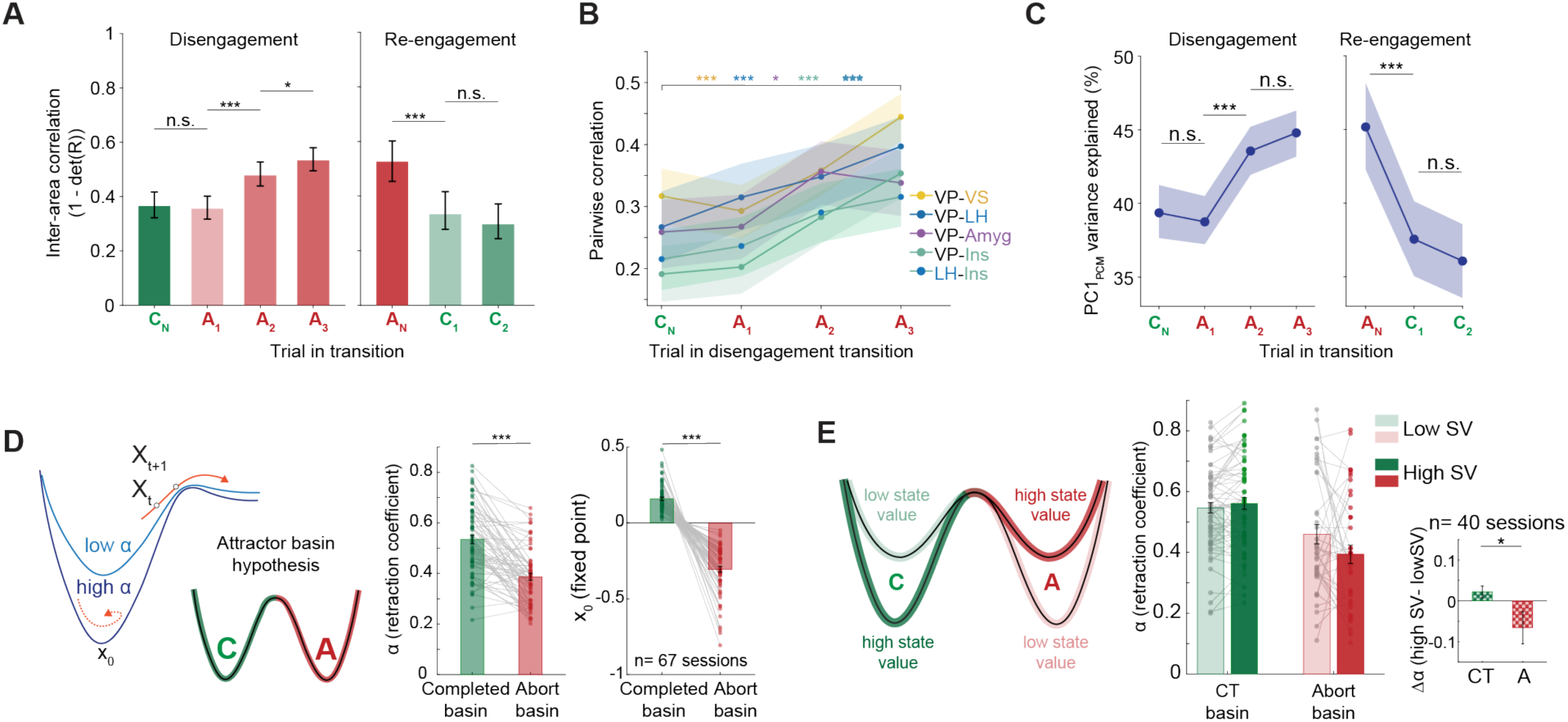
Disengagement is characterized by convergence onto a shared axis with attractor dynamics. **(A)** Inter-area correlation of per-area posterior probabilities across disengagement (left) and re-engagement (right) sequence positions. det(R) refers to the determinant of the correlation matrix, R, of the posterior probabilities from decoding. Data combined across monkeys. Asterisks indicate degree of statistical significance across transitions (Disengagement: C_N_ ◊A_1_, p=0.61; A_1_◊A_2_, p< 0.0001, A_2_◊A_3_, p= 0.026, n=2779 sequences of trials; Re-engagement: A_N_◊C_1_: p = 0.0003, C_1_◊C_2_: p=0.421, n=1093 sequences of trials; error bars from bootstrap procedure). **(B)** Pairwise area correlations across disengagement sequence positions that were statistically significant (C◊A_3_: VP-VS, VP-LH, VP-Ins; p<0.0001; VP-Amyg: p = 0.048; LH-Ins p < 0.0001). **(C)** Variance explained by PC1 of the five-area posterior correlation matrix (PC1_PCM_) during disengagement (left, C_N_ ◊A_1_, p=0.676; A_1_◊A_2_, p< 0.0001) and re-engagement (right, A_N_◊C_1_: p<0.0001, C_1_◊C_2_: p=0.369). **(D)** *Left:* schematic of the attractor basin model hypothesis for engaged (completed trials) and disengaged (abort trial) basins. Low and high retraction coefficients (α) are shown to demonstrate that neural activity at time t (X_t_) is pulled more strongly toward the fixed point (x_0_) when the retraction coefficient is high. *Middle:* Estimated retraction coefficients (α) for engaged and disengaged basins. Both basins had retraction coefficients significantly greater than zero (Completed basin: t(66) =31.277, p<0.0001; Abort basin: t(66) = 28.102, p<0.0001) and significantly less than one (Completed basin: t(66) = -27.208, p<0.0001; Abort basin: t(66) = -44.623, p<0.0001). The engaged basin had a higher retraction coefficient than the disengaged basin (t(66)=8.340, p<0.0001). *Right:* Estimated fixed points (x_0_) for engaged and disengaged basins. The basins had significantly different fixed point positions (t(66)=15.485, p<0.0001). Bars indicate mean parameter values, grey lines connect individual sessions; dots indicate session means (n=67, both monkeys pooled). **(E)** Left: schematic of the hypothesized double dissociation of the effect of state value on basin depths. Right: Attractor fitting split by low and high state value within each basin. The difference in α between high and low state value was larger for the disengaged basin than the engaged basin (paired t-test: t(39) = 2.351, p=0.012, n = 40 sessions; LME SV x basin interaction: t(210)= 2.015, p=0.045, n=67 sessions). Shaded error bars for (A-C) are 95% confidence intervals. (D-E) Error bars: s.e.m.

Both the engaged (C trials) and disengaged (A_2+_ trials) basins had retraction coefficients significantly greater than zero and significantly less than one (Fig. 5D, middle). Values greater than zero confirmed that neural activity was pulled toward the fixed point rather than integrated over time, while values less than one indicated that activity reflected dynamics rather than temporally uncorrelated variability. This demonstrated that population dynamics along the shared axis exhibited attractor-like behavior in both states and ruled out both unconstrained drift and uncorrelated fluctuations around the mean. The engaged basin had a significantly higher retraction coefficient than the disengaged basin (Fig. 5D, middle), indicating that neural activity was more strongly stabilized around the engaged fixed point. The two basins occupied distinct positions in state space (5D, right) and both results held for each monkey individually and survived shuffle testing (fig. S14).

Having established that both states had distinct attractor basins, we then asked whether state value, which modulated disengagement frequency, also modulated basin depth. If state value affected the stability of both motivational states, then the engaged basin would be expected to be deeper when state value is high and shallower when state value is low, and the opposite pattern would be expected for the disengaged basin (Fig. 5E, left). We split completed trials and abort trials within each session into low and high state value groups and fit separate retraction coefficients for each set of trials. High state value was associated with a deeper engaged basin and a shallower disengaged basin, while low state value showed the opposite pattern. These effects were significant across the two basins (Fig. 5E, right). Because expected reward scaled the relative depth of the two basins, the engaged state was the more stable configuration, and more resistant to perturbations when expected reward was higher. Together, these findings demonstrated that state value modulated when disengagement occurred and governed the stability of the neural activity patterns underlying each motivational state.

## Discussion

Our central finding is that motivational disengagement is predicted by the consensus of population activity across limbic areas and that disengagement and engagement correspond to distinct attractor states. Motivational state is not fully captured by expected reward, and our data provide evidence for a motivational signal that operates in parallel to value-based computations. This rests on three empirical pillars. First, neural activity across limbic areas predicts trial engagement before behavior occurs. Second, neural activity in these brain regions converge onto a shared low-dimensional axis as disengagement progresses. Third, the extent of cross-area consensus at disengagement onset predicts whether disengagement will persist. Our results address two gaps in prior work. First, prior accounts of motivation have lacked a continuous readout that dissociates motivation from reward-based value. Second, motivational control has been assigned to individual areas or acknowledged as distributed without a framework for interpreting what that distribution means.

Motivation has resisted precise quantification in part because it is a broad construct. Theoretical work has organized reward-guided behavior around a goal-directed versus habitual distinction (*25–27*) which classifies behavior according to whether it is sensitive to outcome devaluation. That framing captures important variance in how animals execute choices, but a habitual and a goal-directed animal can be either engaged or disengaged. Motivational state operates upstream and modulates the probability of acting. Effort-based measures, including progressive ratio breakpoints and effort discounting (*28–31*), isolate willingness to work for a specific offer. Performance-based measures, such as latency to initiate a trial or time spent exploring an object, capture the frequency of engagement lapses (*32–35*). Trial-by-trial variability in response initiation latency, measured as intrasubject variability in reaction time (ISVRT) in patients, captures the behavioral fluctuations most related to the neural signal we describe here (*36*). None of these approaches, however, characterizes how lapses organize into sequences across trials, which represent a disengaged state, and it is that sequential structure that carries the motivational signal. Performance-based readouts face an additional challenge, in which degraded output can reflect attention lapses, motor errors, or fatigue through mechanisms distinct from motivation. Like task strategy switching (*37–39*), motivation fluctuates across hundreds of trials, and there is structure within disengagement at both behavioral and neural levels that frequency-based and performance-based readouts miss.

Clinical approaches have faced an analogous challenge at a different scale. Dimensional rating scales of anhedonia, anergia, and apathy (*40–43*) are observer-rated or self-reported, and retrospective. The recent shift toward ecological momentary assessments (*44, 45*) reflects the recognition that motivation cannot be captured with static snapshots. Our data support the principle behind this shift: across a two-hour behavioral session, the neural activity moved in and out of engagement on the timescale of seconds, with a neural signature that predicted transitions before they were visible in behavior.

Each area we recorded from is implicated in human motivational and psychiatric disorders. VS and VP are nodes of the striato-pallidal output system implicated in reward and hedonic tone (*12, 46*). The LH has long been tied to homeostatic drive (*47, 48*) and the amygdala and insula carry affective and interoceptive signals (*10, 49, 50*). Prior accounts have assigned motivational function to isolated nodes (*51, 52*). Our results imply that these accounts are not wrong, but they are partial views of a network process. Our data contain a version of this: VP showed the earliest and largest shift along the engagement axis and contributed disproportionately to cross-area coordination. This is consistent with its anatomical position as a convergence point for direct (D1) and indirect (D2) pathways from VS (*53*). It is also consistent with recent work showing suppression of VS projections to VP can reduce trial errors and improves response times (*54*) and direct suppression of VP can increase trial errors (*55, 56*). A single-area account of VP alone would not be wrong, but it would be incomplete.

Human imaging has established that motivation-related signals are distributed across frontal and limbic regions (*57, 58*), and entire functional networks have been associated with motivational readouts using task-based and self-report measures (*59, 60*). Distributed coding has been reported across domains in systems neuroscience (*61–64*). What has remained unclear is what that distribution means computationally: do areas act independently, redundantly, or as part of a collective computation, and how large of a network must be recorded to characterize disengagement?

We propose that a consensus-based system, which we found evidence for in this dataset, provides a path toward interpreting distributed activity. Consensus-based transitions have been described in collective animal behavior, where group decisions emerge from local interactions without any individual having global information (*65*). The same principle has been formalized in distributed computation, where consensus emerges in networks of autonomous agents through local exchange (*66*). One node can dominate the signal, but the collective signal across nodes determines whether a transition occurs. Removing one node could degrade the fidelity of the collective signal without eliminating it. This also accounts for how different single-area perturbations in the brain can produce similar behavioral effects and why manipulating some areas may have no effect at all. Detecting this consensus signal requires resolving population activity at the multi-area circuit level, beyond what mesoscale connectivity measures provide. Consensus across areas could serve as a circuit-level biomarker for motivational state, with implications for closed-loop interventions targeting trial-by-trial variability in engagement, and the sustained motivational deficits observed across psychiatric conditions.

Our consensus result also raises a question: what is the tipping point for motivational transitions? Is a transition related to the number of areas, or the identity of the areas? Causal tests of subsets of areas within the network could elucidate which areas can drive behavioral transitions in disengagement. Prior studies perturbing individual areas in this circuit, including VS and Amygdala in a token-based task, produced inconsistent effects on motivation (*22, 67–74*) and recent stimulation work in human patients only had significant effects on symptoms when VS and VP were co-stimulated (*75*). Under a single-area account, this is puzzling, given the established role of these areas in reward representation, but it is what the consensus framework predicts. Our consensus result therefore predicts that a causal test of motivational disengagement might require simultaneous or combinatorial manipulations of multiple limbic areas, and our framework provides a basis for selecting which subsets to target.

Expanding analyses from the single-neuron level to population-level geometry and dynamics has advanced understanding of how computations are implemented in multiple domains, including decision making (*76, 77*), motor planning (*78, 79*), and working memory (*80, 81*). When we took a similar approach to motivation, we found attractor states for engagement and disengagement that were modulated by expected reward. Basin depth indexes how strongly population activity is held near a state and does not measure the behavioral effort required to maintain that state, but effort may affect basin depth. The engaged basin was deeper than the disengaged basin on average, but expected reward set the relative depth of the two basins.

Attractor basin depth provides a computational analog for motivational states resistant to reversal, as observed in apathy and anergia (*82*), and predicts that chronically reduced reward valuation would deepen disengagement basins and increase resistance to re-engagement. Activity patterns were consistent with persistent activity near each attractor state, though rapid fluctuations between states cannot be ruled out with the current data. Confirming the existence of fixed points and transitions between them will require larger simultaneous datasets and reconstruction of single-trial dynamics. Our characterization of motivation at the level of population geometry and dynamics positions it alongside decision-making and motor planning as computations that operate above the level of specific actions or outcomes.

The population geometry of disengagement supports the hypothesis that motivation is a circuit-level computation. Variance within each limbic area increased while the variance explained by the motivational axis (PC1) also increased during disengagement, meaning the circuit aligned to a shared axis at the expense of each area’s independent functional contribution. This raises the question of whether the same axis would emerge representing engagement and disengagement in the context of other goal-directed behaviors. Past work on feeding, drinking, mating, social and economic decision paradigms have developed rigorous neural accounts that implicate subsets of these same areas (*5, 83–86*). A prediction that arises from our findings is that the shared disengagement axis would re-emerge across limbic areas in other goal-directed tasks with different reinforcers. The conservation of this axis may reflect shared neuromodulatory architecture of these areas across behavioral contexts (*87, 88*). The anatomical areas recruited would likely be conserved, as VP and LH are common substrates across motivated behaviors, while the weighting of individual areas would vary by task. For example, VS showed weaker engagement-related signals in our data, despite being a primary substrate for reward-driven behavior (*89*). The motivational axis we identified was not tied to the specific content of what was being pursued, generalizing across both reinforcer type (tokens versus fluid) and reinforcer identity (water versus mint). An open question remains: is motivation a single, unitary dimension scaled by inputs such as attention, arousal, and effort? Or do those inputs add orthogonal dimensions that expand the dimensionality of motivational state?

Attention, arousal, and effort are also broad constructs that share behavioral signatures with motivational disengagement. These constructs are not equivalent to motivation, and the framework we developed provides a means to dissociate them. Effort costs were matched across trials in our task, and the only penalty for aborting was self-inflicted delay. Despite this, disengagement was structured. Single aborts concentrated at fixation and choice holds, consistent with motor errors, while later aborts were concentrated in not acquiring fixation, suggesting effort avoidance (*58, 90, 91*). First aborts occupied an intermediate position, consistent with a motivational transition rather than a committed shift in state. This structure provides a behavioral scaffold for testing effort dissociations, for example by adding cognitive (*92, 93*) or physical effort (*30, 94*) components to test whether effort costs interact with expected reward to deepen the disengagement basin.

Arousal and attention are harder to dissociate without concurrent physiological measurements. Arousal correlates may covary with the engagement signal (*95–97*), and crossing physiological indices of arousal with the population geometry could specify how much of the motivational axis reflects arousal versus a motivation-specific component. For attention, work has shown that attentional capture by distractors is modulated by current task engagement state and reward (*98*), suggesting the axis we identified may reflect the neural substrate through which motivation gates attentional processing. Individuals with attention deficit disorders can show patterns of disengagement modulated by reward context (*99*), which raises the possibility that some deficits reflect a chronically deepened neural disengagement basin. This is consistent with recent findings that stimulants alter functional connectivity in regions related to arousal and reward, not canonical attention networks (*100*). We predict that the relative contributions of attention and arousal vary across task designs. Accounting for the reward component of the motivational axis provides a reference against which these contributions can be measured and dissociated.

What determines whether an animal engages at all, rather than how it chooses once engaged, is now tractable as a problem of circuit-level computation. Motivation is a network-level variable whose signature is the consensus of population activity across areas. The consensus framework gives distributed activity a computational interpretation and points toward network-level targets for treating motivational deficits in psychiatric disease.

## Acknowledgments

The authors would like to thank J. Napoli, V. Morgan, M. Janssen, A.C. Tangen, K. Guerriero, C. Waters, the Neurophysiology Imaging Facility, and the NIH Animal Care Staff for assistance with monkey handling, surgeries, and animal monitoring; Chipotle, Wittgenstein, and Garbanzo for their behavioral and neural contributions; K. Cameron, C. Rood, P. Pham, and E. Saglio for technical assistance and chamber design; A. Mitz for hardware and software design; V.Costa, S.Shushruth, S.Silas, R.Lam, M.Ramadan, N.Yusif, C. Deister, H. Tang, F. Giarrocco, M. Andujar, A. Mitz, V. Stuphorn, H. Tejeda, S. Lee, B. Ebitz, E. Watts, and D. Pine for project and manuscript draft feedback. This research was supported [in part] by the Intramural Research Program of the National Institutes of Health (NIH). The contributions of the NIH author(s) are considered Works of the United States Government. The findings and conclusions presented in this paper are those of the author(s) and do not necessarily reflect the views of the NIH or the U.S. Department of Health and Human Services.

## Funding

National Institutes of Health ZIA-MH002928(BBA)

## Author contributions

Conceptualization: DCB, BBA

Methodology: DCB, BBA

Formal analysis: DCB, BBA

Investigation: DCB

Resources: BBA

Writing – original draft: DCB

Writing – review & editing: DCB, BBA

Visualization: DCB

Funding acquisition: BBA

## Competing interests

The authors declare that they have no competing interests.

## Data, code, and materials availability

The study involved no preparation of new materials. All data and code used for analyses will be made available upon publication.

## Notes

### Competing Interest Statement

The authors have declared no competing interest.

