## Supplementary Materials for "Consensus across limbic areas predicts motivational disengagement"

**The PDF file includes:**

Materials and Methods
Supplementary Text
Figs. S1 to S14
Tables S1 to S3
References for Supplementary Materials

### Materials and Methods

#### Animal subjects

The experiments were performed on two adult male rhesus macaques (*Macaca mulatta*) that weighed 8-11 kg and were 5-8 years old. The monkeys were pair-housed when possible, and they had access to food 24 hours per day. On testing days, the monkeys were placed on water control and earned fluid through performing the task. Experimental procedures for all monkeys were performed following *the Guide for the Care and Use of Laboratory Animals* and were approved by the National Institute of Mental Health Animal Care and Use Committee.

#### Experimental setup

Monkeys were trained to perform a saccade-based two-armed bandit task. Stimuli were presented on a 19-inch LCD monitor situated 30 cm from the monkeys' eyes. During training and testing, the monkeys sat in a primate chair with their heads restrained. Stimulus presentation and behavioral monitoring were controlled by MonkeyLogic version 2.1 *(101, 102)*. The eye movements were monitored at 400 frames per second using a Viewpoint eye tracker (Arrington Research, Scottsdale, AZ) and sampled at 1 kHz. Fluid rewards were delivered through a mouthpiece with two separate plastic tubes, each connected to a pressurized plastic reservoir gated by a separate solenoid valve. The reservoirs contained either 100% water or mint fluid that consisted of 90% water, 10% Delosi Peppermint Flavor concentrate (Delosi Labs, Kissimmee, FL). The fluids had no caloric differences.

#### Task design

The task was a variant of a token reward reinforcement learning (RL) task *(22, 23)*. In the task, monkeys made choices and received immediate reinforcement in the form of tokens. Each block used four novel images associated with different outcomes including +1 Water token, +2 Water tokens, +1 Mint token, and +2 Mint tokens. The task had six individual conditions defined by the possible values of the image pairs (e.g. +1 Water and +2 Mint). The conditions within a block of 108 trials (6 conditions × 2 counterbalanced for left and right sides × 9 repetitions) were presented pseudorandomly. The monkeys saw each condition twice, once on the left and once on the right, every 12 trials before seeing any condition a third time. Four new images were introduced every 108 trial block. Image sets were novel to the monkeys across blocks and sessions and randomly pulled in sets such that the two monkeys had distinct groups of images. Images provided as choice options were pre-processed and normalized for luminance and spatial frequency using the SHINE toolbox for MATLAB, described previously *(22)*. Monkeys completed 6-11 blocks of trials in a session, with a target of 10 blocks per session.

The monkey began a trial by acquiring fixation at the center of the screen and holding fixation for 500 ms. After a successful fixation hold, two of the four possible images appeared on either side of the fixation spot. The monkey selected a cue by making a saccade to one of the cues. The monkey was required to hold fixation on the chosen image for 500 ms to complete the trial, at which time both choices disappeared and the corresponding number of tokens was added to the corresponding pool of tokens for each of the fluid types. Choosing one of the images led to gaining a corresponding number and kind of token 80% of the time and 20% of the time, the number of tokens did not increase after choice. Mint tokens were shown in light green and water tokens were shown in light blue on the screen above and below the choice targets. The positions of the two groups of tokens were randomized across blocks. At the start of the session, the monkey started with zero tokens. Tokens accumulated across trials and were cashed out for fluids every four to six trials. On cashout trials, following token onset, there was a 1000 ms delay at which time all of the first type of tokens were delivered and subsequently all of the second type of tokens were delivered with 1000 ms delay between fluid types. The order of mint and water delivery was randomized across cashouts. When each drop of fluid was delivered, one token was removed from the screen. Each token was worth approximately 0.22 mL of fluid. All trials, including cashout trials, were followed by a 2000 ms inter-trial interval (ITI), where the monkey was free to move his eyes and accumulated tokens remained visible on the screen.

The monkeys could abort trials in four ways: failure to acquire fixation, failure to hold fixation, failure to select a target, failure to hold a chosen target. The monkey had 2000 ms to acquire fixation once the spot appeared on the screen. If they did not acquire, this was logged as a fixation acquire abort. If the monkey did not hold fixation for 500 ms, this was logged as a fixation hold abort. The monkey had 1000 ms to choose an image and if they did not acquire an image, this was logged as a choice acquire abort. If the monkey chose an image but did not hold fixation on the image for 500 ms, this was logged as a choice hold abort. All aborts were also followed by a 2000 ms ITI before the fixation spot reappeared to start a trial. An aborted trial was repeated until completed, i.e. if the monkey saw the cues appear and aborted the trial, the same cues would appear until the monkey completed the trial.

Surgical procedures

Each monkey was surgically implanted with a titanium headpost. After reaching stable behavioral task performance, a 25 × 35 mm recording chamber was implanted to allow vertical grid access to the Ventral Striatum, Ventral Pallidum, Lateral Hypothalamus, Amygdala and anterior Insula. Procedures followed a previously published approach for semi-chronic implantation of guide tubes for accessing subcortical structures *(23)*. Briefly, chamber placement was planned using pre-operative magnetic resonance imaging (MRI) and was performed without a craniotomy. Following the chamber implantation, the chamber position was verified by MRI with a gadolinium-filled grid inside the chamber. This MRI scan was used to plan recording targets based on individual subject anatomy and atlas coordinates *(103)*. After target planning, small burr holes were drilled above each target area to permit multiple approach trajectories per area, yielding 4-10 usable grid holes per area (i.e. grid holes whose trajectories reached the intended target). For recordings, one MRI-compatible guide tube per area was inserted through a single grid hole, passed through the corresponding burr hole, and lowered 1-4mm below the dura depending on the target. Guide tubes were then secured to the grid and their final positions were confirmed by MRI.

Guide tubes remained in place for 3-20 recording days, during which a multi-contact probe was lowered through each guide tube at the start of every recording session and retracted at the end. Sterile inserts were placed in the guide tubes between sessions. At the end of each 3-20 day period, the full set of guide tubes was removed and replaced. All surgeries and MRI scans were performed under anesthesia. See table S1 for recording session yields.

Neurophysiological recordings

Neurophysiology recordings began after the monkeys had recovered from the chamber implantation surgeries. One 32-channel or 64-channel linear electrode array (V-probe, Plexon Inc, Dallas, TX) was lowered into each guide tube on every recording day. The electrodes had 150-200 µm inter-contact spacing. The probes were advanced to their target location by a micromanipulator (NAN Instruments, Nazareth, Israel) attached to the recording chamber.

Electrophysiological data were acquired with a 512-channel Grapevine System (Ripple, Salt Lake City, UT). The spike acquisition threshold was set at a 4.0 × root mean square (RMS) of the baseline signal for each electrode. Behavioral event markers from MonkeyLogic and eye-tracking signals from Viewpoint were sent to the Ripple acquisition system. The extracellular signals were high-pass filtered (1 kHz cutoff) and digitized at 30 kHz to acquire the single-cell activity. Spikes were manually sorted offline via Plexon Offline Sorter (Plexon Inc, Dallas, TX).

### Behavior analysis

#### *Rescorla-Wagner (RW) model*

To fit choice behavior, a Rescorla-Wagner model *(104)* was fit to the behavioral data. The value update equation was given by the following:

$$v_i(t+1) = v_i(t) + \alpha\left(R_{fluid} - v_i(t)\right) \quad (1)$$

where $v_i$ is the value estimate for cue option *i* that was chosen on trial t, $R_{fluid}$ is the inferred value associated with the change in the number of tokens corresponding to mint or water that followed the choice in trial *t*, and α is the learning rate. The free parameters for reward, $R_{fluid}$ were $R_{mint}$, $R_{+2\ water}$, $R_{+1water}$ in the best fitting RW model. This allowed for the parameterization of the value of mint versus water tokens, which was not known a priori and varied by monkey. The values computed in Eq. 1 were then used to compute choice probabilities for each cue pair using the softmax function:

$$d_j(t) = (1 + e^{(v_i(t) - v_j(t))})^{-1},\ d_i(t) = 1 - d_j(t) \quad (2)$$

where *i* and *j* are the two choice options. Because $R_{fluid}$ was a free parameter, the choice consistency parameter, β, often allowed to be a free parameter in the softmax function, was set to 1. Allowing β to float as well as parameterizing $R_{fluid}$ for all juice levels resulted in an unidentifiable model. The likelihood of the monkey's choices, *D*, given the parameters, was then maximized using the cost function:

$$f\left(D \middle| \alpha, R_{fluid}\right) = \prod_t [\, d_1(t)c_1(t) + d_2(t)c_2(t)] \quad (3)$$

where $d_1(t)$ is the choice probability value for option 1 on trial t, $c_1(t)$ and $c_2(t)$ are indicator variables that take on a value of 1 if the corresponding option was chosen and 0 otherwise. This model was fit across blocks in each session for each monkey to give one set of fit parameters for each session. Only sessions with recording data were used for analyses (Monkey W, n=30 sessions; Monkey G, n=39 sessions).

#### *Characterization of within-condition learning and reaction time drift*

Choice preference and choice reaction times (RTs) were analyzed as a function of trial position within each of the six cue conditions across sessions. Each monkey's behavior was fit separately with a RW-RL model. Data from both monkeys was then concatenated and analyzed jointly for behavioral summaries. For each condition and trial position, mean and standard error were computed across sessions for behavioral choices and RW-RL choice probability.

To assess choice preference, a one-sample t-test was performed across sessions testing whether the mean choice frequency exceeded chance. Resulting p-values were adjusted for multiple comparisons using the Benjamini-Hochberg false discovery rate procedure (BH-FDR, α=0.05). To characterize within-condition RT drift, a log-linear drift model was fit to per-session RTs. For each session and condition, RT as a function of trial position was modeled as:

$$d_j(t) = \beta_o + \beta_{drift} * t + \beta_{early} * \log(t) \tag{4}$$

where $\beta_{drift}$ captures the linear accumulation of RT across trials and $\beta_{early}$ accounts for the early RT jump in the first trials of a block. The distribution of session-level $\beta_{drift}$ estimates was tested using a one-sample, right-tailed t-test for positive drift with correction for multiple comparisons across conditions.

*State-value Markov Decision Process (MDP) model*

State values were computed using a Markov decision process (MDP) model adapted from a previously published model *(21)*. The structure of the model and the modifications required for the current task are described below.

The MDP model computes the value of each task state, defined as the sum of immediate rewards and the expected discounted sum of future rewards from that state. This is referred to throughout as the state value (SV) or the utility of the state $u(s_t)$. The state $s_t$ is a function of five task features: the number of accumulated water tokens ($NTk_{Water}$) and mint tokens ($NTk_{Mint}$), the number of trials since cash-out (TSCO), the task epoch (TE), and the number of observations of a cue pair within a block (NObs). All pairwise combinations of ($NTk_{Water}$, $NTk_{Mint}$) satisfying $NTk_{Water} + NTk_{Mint} \leq 12$ were included as valid states, with a maximum of 12 total tokens consistent with the maximum of six trials before cashout. The state space consisted of all valid combinations of these five features. The state value was computed using value iteration *(105)* applied to the following Bellman optimality equation:

$$u(s_t) = \max_{a_t \in A} \left[ r(s_t, a_t) + \gamma \sum_{j \in S_{t+1}} p(j|s_t, a_t) u(j) \right] \tag{5}$$

where $s_t$ is the state, $a_t$ is the action taken at that state, $u(s_t)$ is the state value, $r(s_t, a_t)$ is the immediate reward, γ is the discount factor, $p(j|s_t, a_t)$ is the transition probability to future state $j$ and $S_{t+1}$ is the set of immediate future possible states from state $s_t$ if one takes action $a_t$. The summation over future states ($j$) is the expected future utility, taken across the transition probability distribution $p(j|s_t, a_t)$. As in the original published model, the discount factor was γ=0.999. Value iteration was run until the policy and state values converged, which required approximately 100 iterations.

The original model was designed for a task with a single token type and included token count as a discrete scalar dimension of the state (NTk, range 0-12). The present task used two distinct token types (water and mint), requiring the token-count dimension to be expanded into two separate state features, $NTk_{Water}$ and $NTk_{Mint}$. All other state features and transitions were identical to those in the published MDP model for tokens *(21)*. The transition probabilities from fixation to any of the six cue states were modeled as p(cues) = 1/6. Cash-out transition probabilities were p(cashout) = 0 for TSCO 1–3, p(cashout) = 0.33 for TSCO = 4, p(cashout) = 0.50 for TSCO = 5, and p(cashout) = 1.0 for TSCO = 6. As in the original model, transition probabilities for the token outcome given a choice (p(ΔNTk | choice)) were derived from the RW model fits to each monkey's choice behavior (see RW model section), following the same parameterization previously described *(21)*.

In the original task, tokens of a single type were each worth one unit of juice at cash-out, so the immediate reward at cash-out was simply the total token count. In the task used in this study, water and mint tokens were redeemed sequentially at cash-out but carried different subjective values. Because the relative value of each token type was not known a priori, the cashout reward used in the MDP was not the raw token count but a weighted sum of water and mint tokens, with weights derived from the RW model fitted to each monkey's choice behavior. The RW model yielded free parameters representing the subjective values of each reward magnitude: $R_{Mint}$, $R_{+1Water}$, and $R_{+2Water}$ (see Rescorla-Wagner (RW) model). The ratio of $R_{Mint}$ to $R_{+1Water}$ provided an empirical estimate of how one mint token was valued relative to one water token for each monkey. This ratio was used to scale mint token counts in the cash-out reward. The total immediate reward delivered at cash-out was therefore:

$$r_{cashout} = NTK_{Water} + \lambda * NTK_{Mint} \tag{6}$$

where $\lambda$ is the monkey-specific mint scaling factor derived from the RW parameter estimates ($\lambda$=0.1 for Monkey W; $\lambda$=0.7 for Monkey G). This formulation ensured that the state value reflected each monkey's learned relative preference for the two token types, rather than treating them as interchangeable. This enabled the choice-probability curves derived from the MDP to match those derived from the RW model. A separate MDP was fit for each monkey using its corresponding λ and transition probabilities using the larger state space that accounted for both token types.

State values were extracted for all trials and epochs using the MDP fits for each monkey. This produced a table of states such that the value of each state was: $V_{s_t} = u(s_t) = f(NTkWater, NTKMint, TSCO, TE, NObs)$. These state values were used to characterize trial-by-trial relationships to reaction time to acquire fixation, choice reaction time, and trial aborts.

Mean reaction times were computed by averaging reaction times across blocks of trials and then averaging across sessions for each monkey. Outlier RTs were removed using Tukey's method: RT > q0.75+ 1.5*IQR and RT< q0.25 – 1.5*IQR, where IQR is the interquartile range within session.

To assess the relationship between state value and reaction times to acquire fixation, the following linear regression on fixation state value was computed:

$$RT_{acquire\_fixation} = \beta_0 + \beta_{V_{Fix}} V_{fix} \tag{7}$$

where $V_{fix}$ is the value of the fixation state in the corresponding trial.

To assess the relationship between state value and choice reaction times, the following linear regression was computed:

$$RT_{choice\_reaction\_time} = \beta_0 + \beta_{V_{Cue}} V_{cue} + \beta_{\Delta V}(\Delta V) \tag{8}$$

where $V_{cue}$ is the state value at cue onset and $\Delta V = V_{cue} - V_{fix}$. The variable $\Delta V$ captures the change in value when the cues are revealed.

To assess the relationship between state value and the probability of aborting a trial, the following logistic regression was computed:

$$p(Abort) = (1 + e^{-(\beta_0 + \beta_{V_{cue}} V_{cue}} )^{-1} \tag{9}$$

where $V_{cue}$ is the value of the cue presentation state. To assess whether regressors were significantly different than zero, for each monkey, t-tests on the distributions of beta values across sessions for each regressor were performed for each monkey.

To visualize the probability of abort predicted by the model versus the frequency of abort at different state value levels, trial-level abort probabilities were predicted for each trial using the session-specific logistic regression parameters and each trial's $V_{cue}$ values. Predicted probabilities and actual abort outcomes were then pooled across sessions for each monkey and binned by $V_{cue}$ in bins. Mean predicted abort probability and actual abort frequency were computed within each bin.

*Abort patterns and sequences*

To characterize the temporal dynamics of motivational disengagement and re-engagement, abort sequences were defined as runs of one or more consecutively aborted trials (referred to as $A_i$ where *i* indexes the ordinal abort trial) bounded by completed trials (referred to as $C_i$). A sequence began on the trial immediately following a completed trial and ended on the last aborted trial before the next completed trial. The terminating completed trial was not counted as a member of the sequence. Within each sequence, individual aborts were indexed $A_1$, $A_2$, ..., $A_N$, where $A_1$ is the first abort following the preceding completed trial and $A_N$ is the last abort immediately preceding the next completed trial. The preceding completed trial before an abort sequence is denoted $C_N$, and the first through fourth completed trials following the sequence are denoted $C_1$ through $C_4$.

To characterize how abort rate evolved across a session a per-trial binary abort indicator (1 = abort, 0 = completed) was computed from the abort codes (for fixation acquisition, fixation hold, choice acquisition, choice hold) from MonkeyLogic. Sessions were NaN-padded to the maximum trial count observed across all sessions, and mean abort rate and SEM were computed across sessions at each trial position, including only positions at which at least 12 sessions contributed data. The resulting mean and SEM traces were smoothed with a Gaussian kernel (window = 30 trials) for visualization.

To characterize the distribution of abort sequence lengths, abort sequences were identified within each session as runs of consecutive abort trials. For each pooled length bin (1, 2, 3, 4, 5, 6–10, 11–15, 16–20, and ≥21 consecutive aborts), the mean number of sequences per session was computed across sessions.

To examine how abort sequence length varied with session progress, each trial's position within a session was normalized by the total trial count for that session, yielding a progress index on [0, 1]. Sequences were binned by their session progress position and by their length, and the mean fraction of sequences in each length category was computed per bin across sessions.

For behavioral analyses of re-engagement, sequences were required to contain at least three consecutive aborts ($N \geq 3$) to ensure sufficient preceding disengagement context. Completed trials following qualifying sequences were extracted and indexed $C_1$ through $C_4$ relative to the sequence end.

For all neural analyses, sequences were selected to ensure sufficient context on both sides of the behavioral transition of interest. Analyses of disengagement required a minimum of three preceding completed trials and three consecutive aborts ($\ldots C_{N-2}C_{N-1}C_NA_1A_2A_3\ldots$), and analyses of re-engagement required a minimum of three consecutive aborts followed by three completed trials ($\ldots A_{N-2}A_{N-1}A_NC_1C_2C_3$). These criteria did not change even if only the positions immediately surrounding the transition were analyzed (the abort-to-completed boundary for re-engagement, and the completed-to-abort boundary for disengagement) or presented in figures. Sessions were included in neural analyses only if they contributed at least five qualifying sequences.

Neural analysis

*Drift correction*

Population spiking activity exhibits slow, session-timescale fluctuations that can be either due to electrode drift or a slow varying signal that is correlated with elapsed time within a recording session *(97)*. Because abort frequency increased across sessions, a neural signal tracking session time could artifactually appear to discriminate upcoming trial engagement (C or A). To remove this confound prior to all analyses, dimensionality reduction was performed on the trial-mean population activity during the ITI within each session, independently per area and cell, and the first principal component ($PC1_{DC}$, where DC = Drift Correction) was subtracted from all downstream activity.

For each session, each unit's mean firing rate (Hz) across the pre-fixation ITI window (−2000 to 0 ms relative to fixation onset) was computed to produce a matrix of [trials × units]. This matrix was z-scored per unit using each unit's within-session mean and standard deviation, then used in PCA without trial-type labels. The $PC1_{DC}$ loading vector, estimated from the trial-mean data, was used to remove drift from the trial-by-trial mean rates and trial-by-trial binned firing rates used in downstream analyses. For each time bin within the ITI, the trial's population activity vector was projected onto $PC1_{DC}$ and the resulting scalar projection was subtracted from the population activity vector for that trial and bin. This operation removed the component of the firing rate in each bin that lies along the session drift axis, while leaving all orthogonal structure intact.

The adequacy of drift correction was evaluated using the regression framework described in *Regression of Posterior Probabilities on State Value and Time in Session*. Briefly, posterior probabilities from decoding were regressed within each session on two z-scored regressors: the MDP-derived state value and a satiety proxy of time-in-session. Resulting beta coefficients were tested across sessions. Prior to any correction, PC1, referred to as $PC1_{DC}$, of the trial-mean population activity accounted for approximately 80% of the within-session correlation between the posterior probabilities and time. After $PC1_{DC}$ subtraction, the residual beta coefficients for time-in-session were reduced to account for 0-10% of the within-session correlations. The remaining betas displayed a pattern inconsistent with a global recording artifact, as the residual correlation varied in sign across areas and trial types and was asymmetric across monkeys. The residual correlations were also resistant to two additional corrections, removing $PC1_{DC}$ and $PC2_{DC}$, and a targeted dimensionality reduction method. First, $PC2_{DC}$ of the trial-mean population activity was subtracted in addition to $PC1_{DC}$. Second, a targeted dimensionality reduction was applied in which each neuron's $PC1_{DC}$ residuals were regressed against true elapsed session time, a population-level unit vector was constructed from those regression weights, and its projection was subtracted from all binned activity. Neither approach produced a consistent or meaningful reduction in the residual time-in-session beta coefficients beyond what $PC1_{DC}$ removal alone achieved. These null results supported limiting correction to $PC1_{DC}$.

After $PC1_{DC}$ subtraction, residual firing rates were z-scored across all trials within each session and mean-centered. The resulting residuals, stored in 50, 200, or 500 ms bins, served as input to other analyses.

*Responses to task features*

To identify neural responses to different task components, a sliding window multi-way ANOVA model was fit to spike counts computed in 200 ms bins, advanced in 50 ms increments. ANOVA models were fit to capture the entire trial time, each locked to fixation onset, cue onset, and token outcome to account for fixation and choice reaction times which varied by trial. Factors included the number of tokens of each type on the screen (#token of Mint, Water), the change of token numbers (Δtokens of Mint, Water), the drops of fluid delivered at juice outcome period (#Mint, # Water), assigned a priori value (+2, +1), RW cue value, MDP state value, block number, chosen stimulus identity, and chosen direction (left or right). The stimulus identity was the specific image used in each block to represent each outcome, which was the interaction of chosen value and block number. The factor block number was used to account for non-stationarity due to drift.

*Responses to task features during the ITI*

To identify neural responses to task features relevant only during the ITI, a separate ANOVA was constructed. This ANOVA was run on the mean drift-corrected activity during the ITI and binned drift-corrected activity across the ITI (200 ms bins, 50 ms sliding). The eight factors included three task events from the previous trial that were relevant to the ITI: change of Mint tokens (Δ Tokens Mint), change of Water tokens (Δ Tokens Water), and a categorical variable as to whether the previous trial was a cashout trial. The other five factors included were the # of mint tokens, the block number in the session (to account for drift), the trial number in the block (to account for learning), MDP state value, and upcoming trial outcome (Completed or Abort). Each unit was tested for significant coding of each of the eight factors ($p < 0.05$).

For each area, units were cross-classified by whether they showed significant encoding ($p < 0.05$) of state value and of upcoming trial outcome, and the resulting 2x2 contingency table was tested against the null of independence using a Pearson chi-square test (1 df). Enrichment was assessed by comparing the observed number of co-encoding units to the count expected under independence, given by the product of the two marginal proportions and the total unit count. Tests were performed separately for each of the five areas, and significance was evaluated against a Bonferroni-corrected threshold of $p < 0.01$ (0.05 across five areas).

*Dimensionality reduction using principal components analysis (PCA)*

PCA was applied at multiple stages of the neural analysis pipeline. The first application, described in detail in *Drift correction*, was performed on trial-mean population activity (one mean firing rate per ITI per unit) to identify and remove the dominant session-timescale drift component prior to all subsequent analyses. This was done on the activity averaged over the 2000 ms ITI. The second application was performed on the drift-corrected, residual mean rates to reduce the dimensionality of the population activity for visualization and distance metrics. For this second round of PCA, firing rates from all ITI 200 ms time bins were concatenated across trials within each session, z-scored per unit, and submitted to PCA separately per area and monkey. The resulting low-dimensional projections captured the dominant axes of across-trial and across-time variance in the drift-corrected activity and were used to visualize population activity in state space and compute Euclidean distances in PC space (see Euclidian distance measures). The same dimensionality reduction steps were used for binned ITI activity to characterize attractor dynamics (see Attractor dynamics analysis). The number of PCs retained for each analysis is specified in the corresponding sections. The components from dimensionality reduction on the drift-corrected activity for visualization are noted as PC#. The components for dimensionality reduction on the posterior correlation matrix are noted as $PC\#_{PCM}$ (see *Posterior coherence analysis*).

*Decoding from ITI activity*

Motivational state (completed vs. abort trial) was decoded from single-session population activity using a linear support vector machine (SVM) classifier implemented in MATLAB. The same procedure was applied to the mean residual ITI activity after removing the PC1 component and binned residual ITI activity after removing PC1. For each session, the feature matrix consisted of either trial-mean firing rates from the ITI (2000 ms) or binned rates (10 200 ms bins) across all recorded units in a given area, with the predictor variable being the binary trial outcome label (completed or abort) assigned to each ITI. To address class imbalance, which was substantial across sessions given that abort trials were less frequent than completed trials in most sessions, inverse-frequency sample weights were assigned to each trial prior to fitting ($w = 0.5/n_{Class}$), such that each class contributed equally to the SVM objective regardless of its raw count.

Classification performance was estimated using stratified k-fold cross-validation ($k = 10$), preserving the class ratio within each fold. Accuracy was defined as the fraction of correctly classified out-of-fold trials. To establish a null distribution, the full cross-validated decoding procedure was repeated on a single permutation of the trial labels for each session, yielding a shuffle accuracy for comparison. Decoding accuracy was evaluated against shuffle at the population level across sessions using a one-sided paired t-test.

*Computation of posterior probabilities from decoding*

The raw output of each cross-validated SVM fold is a scalar decision score reflecting the signed distance of each test trial from the learned hyperplane, where positive values favor the abort class and negative values favor the completed trial class. These scores are not calibrated probabilities. Because class frequencies were imbalanced across sessions, direct conversion via standard Platt scaling *(106)* as implemented in MATLAB's fitPosterior was inadequate, as this approach fits a single sigmoid to the full training set without accounting for sample weights and produces biased probability estimates under unequal class frequencies.

To obtain well-calibrated posterior probabilities under class imbalance, a weighted Platt scaling procedure was applied within each cross-validation fold *(107)*. For each fold, a logistic regression model was fit to the out-of-fold decision scores from the training partition using class-balanced frequency weights. For each training trial, an inverse-frequency class weight was first computed as $w_i = 0.5 / n_{Class}$, where $n_{Class}$ is the number of trials belonging to that trial's class within the training fold. This ensured each class contributed equally to the fit regardless of underlying trial count. These weights were then multiplied by the total number of training trials in that fold ($w_i * n_{Train}$) to convert them from a relative importance scalar to an effective trial-replication count. This made the weights compatible with MATLAB's glmfit frequency-weight convention, in which the weight is interpreted as a trial replication count rather than a per-trial importance scalar. The fitted sigmoid was then applied to the held-out test-fold decision scores, yielding a calibrated estimate of P(abort | x) for each test trial. This procedure was repeated independently for each cross-validation fold. No trial's posterior was computed from a fold in which that trial contributed to classifier training.

Two posterior quantities were retained for each trial. P(true class | x) is the calibrated probability that the upcoming trial after the ITI was the trial label, and was defined as P(abort | x) for abort trials and 1 - P(abort | x) for completed trials, reflecting the classifier's confidence in the correct assignment for each trial.

*Regression of posterior probabilities on state value and time in session*

To assess the independent contributions of motivational state and time-in-session to the decoded neural state, the cross-validated posterior probability of the true class, P(abort | x) or P(completed | x), was regressed against two predictors within each session for each area separately. The two regressors for a single model were MDP-derived state value (SV) and a time-in-session metric that served as an estimate of satiety (SAT). Regressions were fit separately for completed trials and abort trials. Both predictors and the dependent variable were z-scored within session prior to fitting, yielding semi-standardized regression coefficients interpretable as the change in posterior probability (in standard deviation units) for a one standard deviation change in each predictor.

Time-in-session was operationalized using four metrics: (1) trial count including aborts, (2) trial count excluding aborts (i.e., advancing only on completed trials), (3) elapsed time in session in seconds, and (4) cumulative fluid (mL) received in session. Each was computed from the behavioral record and z-scored within session before entry into the regression.

Three model families were fit for each session, condition, and area. The full model took the form:

$$\hat{P}_t = \beta_0 + \beta_{SV} SV + \beta_{SAT} SAT_t + \varepsilon_t \quad (10)$$

where $\hat{P}_t$ is the z-scored posterior probability for trial t, SV is the z-scored MDP state value from the ITI epoch, SAT is the z-scored session-progression metric at trial t, and $\varepsilon$ is the residual. Nested models omitting one predictor were also fit: an SV-only model ($\beta_{SAT}$=0) and a SAT-only model $\beta_{SV}$= 0. The full model was fit once for each of the four SAT operationalizations, yielding four candidate full models per session. All models were fit using ordinary least squares.

Model selection across the four full model variants and the SV-only model was performed using the Bayesian Information Criterion (BIC), averaged across sessions within each area and condition. The full model variant with the lowest mean BIC was selected as the best full model. Regression coefficients from the best full model were pooled across sessions, and significance was assessed using a one-sample t-test against zero across sessions.

*Computation of changes in posteriors across disengagement and re-engagement*

To characterize the dynamics of the decoded motivational state across disengagement and re-engagement transitions, the progression of per-trial posterior probabilities was examined relative to the position of each trial within a qualifying behavioral sequence. For each transition epoch, a single slope value was computed per session by pooling all eligible trials across qualifying sequences within one session and fitting an ordinary least squares regression of posterior probability on trial position. This yielded one slope per session per area, which was then tested across sessions.

For disengagement, two slopes were computed. The pre-abort slope across completed trials was fit using the last three completed trials of each qualifying run ($C_{N-2}$, $C_{N-1}$,$C_N$), with trial position coded as ordinal distance from the end of the run, and tested against the one-tailed alternative that the slope was negative (i.e., posteriors declining toward the transition). The abort-run slope was fit using the first four abort trials of each qualifying abort run ($A_1$ - $A_4$), with abort ordinal as the predictor, and tested against the one-tailed alternative that the slope was positive (i.e., posteriors rising across the abort sequence). Sessions contributing fewer than five qualifying transitions were excluded.

For re-engagement, two slopes were computed across the abort-to-completed transition. The pre-completed trial slope was fit using abort trials falling in the latter portion of qualifying abort runs ($A_{N-2}$, $A_{N-1}$,$A_N$), with abort position as the predictor, and tested against the one-tailed alternative that the slope was negative. The completed-run slope was fit using the first three completed trials following the transition ($C_1$, $C_2$, $C_3$), and tested against the one-tailed alternative that the slope was positive.

All slope tests were conducted separately for each monkey and on sessions pooled across both monkeys. Per-area significance was assessed using t-tests against zero, with Holm-Bonferroni correction applied across the five areas. Differences in slope magnitude across areas were assessed using a one-way ANOVA on the pooled session-level slopes. Differences between areas (VP > VS and LH > VS) were assessed with one-tailed t-tests corrected for multiple comparisons.

*Euclidian distance of first-abort ITI activity from the completed trial mean*

To test whether the neural state at the first abort of a sequence ($A_1$) predicted the subsequent depth of disengagement, the distance of each $A_1$ trial's population activity from the mean completed-trial population state was computed. This analysis was carried out in four variants to assess robustness of the result: (1) projection onto the first three PCs of the binned residual activity after PC1 removal from mean activity (presented in Results) (2) projection onto the session-specific SVM decoding axis using trial-averaged, binned activity across the ITI window, (3) projection onto the session-specific SVM decoding axis using mean ITI activity, and (4) analyses conducted separately for each monkey rather than pooled. All four approaches yielded qualitatively identical results, with $A_1$ distance increasing monotonically across sequence-length categories in all areas.

For the approach presented in the main text, dimensionality reduction was conducted a second time on the drift-corrected ITI activity for each area. Each trial was then projected onto the first 3 principal components. The completed trial mean ($\mu_{CT}$) was computed across all completed trials in the session, and the distance for each $A_1$ trial was defined as the deviation between the single trial projection and $\mu_{CT}$, normalized by the standard deviation of completed trial activity within that session ($\sigma_{CT}$):

$$d_{A1} = \frac{proj_{A1} - \mu_{CT}}{\sigma_{CT}} \tag{11}$$

Normalization by $\sigma_{CT}$ was applied to place distances on a common scale across sessions and monkeys, since the absolute magnitude of the projections varied with the number of recorded units and the geometry of the lower dimensional state space in each session.

$A_1$ trials were identified as abort trials immediately preceded by at least three consecutive completed trials, to ensure a stable baseline engagement state prior to the first abort. Each $A_1$ trial was then categorized by the length of the abort sequence it initiated: category 1 ($A_1C_1$, isolated abort followed immediately by a CT), category 2 ($A_1A$, one additional abort), category 3 ($A_1AA$ and $A_1AAA$, two or three additional aborts, pooled to ensure sufficient trial counts per session), and category 4 ($A_1AAAA+$, four or more additional aborts). Sessions were included only if all four categories contained at least five $A_1$ trials; sessions not meeting this criterion were excluded. The session-mean distance was computed within each category, and grand means and standard errors were computed across sessions pooled from both monkeys.

Statistical significance of the difference in $A_1$ distance between category 1 and category 4 was assessed using a paired t-test across sessions, with Benjamini-Hochberg false discovery rate correction applied across the five areas.

*Consensus analysis using Euclidian distance*

The $A_1$ distance analysis was extended to predict next-trial outcome from the joint pattern of $A_1$ displacement across areas on single trials. The normalized $A_1$ distances computed in the preceding analysis were retained. Each $A_1$ trial was labeled by the identity of the trial immediately following ($A_2$ if the next trial was an abort, $C_1$ if it was a completed trial). All $A_1$

trials from sessions with at least three of the five brain areas simultaneously recorded were included in this analysis (N = 65 sessions: 11 sessions with 3 areas, 18 with 4 areas, 36 with 5 areas; n = 11,989 $A_1$ trials pooled across both monkeys).

For each session and each recorded area, a per-area threshold τ was defined as the median $A_1$ distance within that session and area. Each area was then assigned a binary vote on each $A_1$ trial: 1 if the trial's normalized distance exceeded τ and 0 otherwise. By construction, this yielded a marginal vote rate of 0.5 within each session and area, removing differences in absolute distance scale across sessions and monkeys as a contributor to vote rate. The vote fraction on each trial was defined as the number of areas voting 1 on that trial out of the number of recorded areas.

The relationship between vote fraction and the immediately following outcome ($A_2$ versus $C_1$) on trial *i* in session *j* was modeled with a generalized linear mixed-effects model (GLMM) of the form:

$$logit\left[\, p(next = A_2)_{ij}\right] = \beta_0 + \beta_{VF} VoteFraction_{ij} + \beta_m 1\left( monkey_j = W \right) + u_j \qquad (12)$$

where $\beta_0$ is the fixed intercept, $\beta_{VF}$ is the fixed effect of vote fraction, $\beta_m$ is the fixed effect of monkey identity, and $u_j$ is the random intercept for session *j*. The model was fit by Laplace approximation in MATLAB (fitglme). Significance was assessed via Wald test on $\beta_{VF}$. 95% confidence intervals on the empirical P(next=$A_2$) were computed using the Wilson score interval. Confidence intervals on the model-predicted P(next=$A_2$) at each vote fraction level were obtained by parametric bootstrap (2,000 iterations). Samples were drawn from the multivariate normal distribution with mean equal to the fitted coefficients and covariance equal to the coefficient covariance matrix returned by fitglme.

Three control analyses were performed to confirm that the observed relationship between vote fraction and outcome reflected trial-level cross-area coupling rather than properties of any single area or chance alignment of vote fraction with outcome. All three controls were performed separately for sessions stratified by number of recorded areas (3-, 4-, 5-area subsets).

The first control tested whether area votes were positively coupled across trials more than would be expected if each area voted independently. Because the median threshold yields a marginal vote rate of 0.5 within each session and area by construction, the independence null is that vote count on each trial follows Binomial($N_{areas}$, 0.5). The observed vote count distribution at each area-set size was compared to the binomial null using a Pearson chi-square goodness-of-fit test.

The second control tested whether the consensus relationship required trial-level coupling between areas rather than each area's marginal vote rate alone. Within each session and area, $A_1$ distances were permuted across trials, destroying trial-level coupling between areas while preserving each area's marginal distribution of distances within session and trial count structure. The per-session per-area median threshold was recomputed on the permuted distances, votes and vote fraction were recomputed, and the consensus mixed-effects model was refit. This procedure was repeated 1,000 times to construct a null distribution of $\beta_{VF}$. Significance was assessed using the one-tailed permutation p-value defined as the proportion of permuted $\beta_{VF}$ values greater than or equal to the observed $\beta_{VF}$.

The third control tested whether the vote fraction effect survived shuffling that broke the trial-level link between vote fraction and outcome. Within each session, next trial outcomes ($A_2$ or $C_1$) were permuted across $A_1$ trials, leaving the observed vote fraction on each trial unchanged.

The consensus mixed-effects model was refit on each permuted dataset, and this procedure was repeated 1000 times to construct a null distribution of $\beta_{VF}$. Significance was assessed using the same method as the first control analysis.

The primary analysis used the per-session per-area median threshold. Three fixed thresholds ($\tau$ = 0.5, 1.0, and 1.5 $\sigma_{CT}$ units) were also tested as sensitivity analyses. The direction and significance of the consensus relationship were preserved across all four threshold choices, but the median threshold gave the most consistent results across sessions and monkeys, because absolute distance scales varied modestly between sessions and monkeys despite the within-session $\sigma_{CT}$ normalization.

*Computation of mean rates over disengagement and re-engagement transitions*

Mean z-scored ITI activity was plotted across ordered trial positions spanning the disengagement ($C_1$–$C_2$–$C_3$–$A_1$–$A_2$–$A_3$–$A_4$) and re-engagement ($A_1$–$A_2$–$A_3$–$C_1$–$C_2$–$C_3$) sequences for units that were identified as significantly responsive to abort versus completed-trial ITIs. Sessions were included only when they had at least five qualifying sequences. In both analyses, trial-level residuals within each position bin were averaged per session to produce one trace per session per area, and session-level traces were then averaged across sessions. Data from both monkeys were pooled. For visualization, the concatenated session-mean trace was smoothed using a Gaussian kernel (window of 8 bins at 200 ms per bin, spanning approximately one full ITI) applied across the full concatenated sequence. All statistical comparisons were performed on unsmoothed data. Significance of transitions was assessed by computing the mean activity over a 2-second window at the end of the reference position ($C_3$ for disengagement; $A_3$ for re-engagement) and at the start of each subsequent position, taking the pairwise difference per session, and testing each difference against zero using a one-sample t-test. Holm-Bonferroni correction was applied across the set of comparisons within each area (4 comparisons for disengagement, 3 for re-engagement).

*Posterior coherence analysis*

To characterize the joint structure of the decoded motivational state across limbic areas during progressive disengagement and re-engagement, the cross-validated posterior probability of abort, P(abort | x), was extracted for each trial and area as described in *Computation of posterior probabilities from decoding*. For each qualifying disengagement transition $CA_1A_2A_3$ and re-engagement transition $AC_1C_2C_3$, the posterior probability from each area at each step-bin was extracted and assembled into a pooled observation matrix of dimension $N \times 5$, where N is the total number of qualifying sequences across sessions and monkeys, and the five columns correspond to the five recorded areas (VS, VP, LH, Amygdala, Insula). Sessions with fewer than five qualifying sequences (CCCAAA or AAACCC) were excluded. Because not all five areas were simultaneously recorded in every session, all matrix operations used pairwise-complete observations.

For each step-bin, a $5 \times 5$ Pearson correlation matrix R was computed from the pooled observations. The determinant of R, det(R), bounded in [0, 1], served as the primary measure of inter-area posterior coherence: det(R) = 1 when all areas are uncorrelated and approaches 0 as inter-area correlations increase. Because det(R) is computed on the correlation matrix rather than the covariance matrix, it reflects the correlation structure of the posteriors independent of each area's marginal variance. The variance of each area's posterior across the qualifying sequences was separately quantified as the diagonal entries of the covariance matrix computed from the same pooled observations, providing a measure of how much each area's motivational state representation fluctuated across sequences within each step-bin. Principal components analysis was applied to each step-bin's correlation matrix (R) and the fraction of variance explained by PC1 (labeled as $PC1_{PCM}$ for Posterior Correlation Matrix for distinguishing it from other PC1s) was retained as a measure of the dimensionality of the joint posterior distribution. An increase in $PC1_{PCM}$ explained variance indicated that inter-area posterior covariation was concentrated along a single shared axis.

Confidence intervals were obtained by paired bootstrap resampling (1000 iterations) of the pooled sequence observations. In each iteration, a single set of N row indices was drawn with replacement from the N qualifying sequences pooled across all sessions and both monkeys, and the same indices were applied to all four step-bin matrices simultaneously, preserving the pairing of observations from the same behavioral sequence. For each iteration, det(R) was recomputed at each step-bin, and the difference in det(R) between consecutive step-bins was recorded, yielding a bootstrap distribution of 1000 difference values per consecutive pair. For each consecutive step comparison, the observed difference in det(R) was compared against a bootstrap distribution of 1000 resampled differences. The one-tailed p-value was defined as the proportion of bootstrap differences inconsistent with the predicted direction of change, and Holm-Bonferroni correction was applied across the three consecutive step comparisons within each transition direction. All analyses were conducted on data pooled across both monkeys and replicated separately for each monkey.

The same paired bootstrap procedure was used to test two additional quantities derived from each step-bin's pooled observations. First, the per-area variance, quantified as the diagonal entry of the covariance matrix for each area, was tested for a monotonic change from the first to the last step-bin of each transition ($C_N$ to $A_3$ for disengagement; $A_N$ to $C_3$ for re-engagement). For each area, the observed difference in variance between the terminal and initial step-bins was compared against the bootstrap distribution of that difference, yielding a p-value; Holm-Bonferroni correction was applied across the five areas. Second, the fraction of variance explained by PC1 of the correlation matrix ($PC1_{PCM}$) was tested for consecutive step changes using the same bootstrap proportion test and the same correction across the three consecutive step pairs.

To characterize how different limbic areas loaded onto $PC1_{PCM}$, a separate analysis pooled all completed trials and all abort trials excluding first aborts (i.e. no $A_1$) into two groups. The correlation matrices and dimensionality reduction steps were then repeated to produce the loadings.

<u>*Attractor dynamics analysis*</u>

Attractor dynamics analyses were conducted using a previously published linear dynamical systems framework *(24)*. The structure of the model and the modifications required for the data in this manuscript are described below.

Neural population activity was projected onto the first principal component of a grand cross-area basis computed across all simultaneously recorded units within each session after drift correction (see <u>*Dimensionality reduction using principal components analysis (PCA)*</u>). This yielded a scalar time series $X_t$ and $X_{t+1}$ for each trial and 200 ms time bin. Bases were computed across all areas jointly to maximize statistical power but separately by monkey. All subsequent attractor fitting was performed on these 1-D projections (200 ms bins, 10 bins per ITI) for each session.

Following previous work *(24),* the bin-to-bin evolution of population activity was modeled as a first-order, undriven linear system with a fixed point:

$$X_{t+1} = X_t - \alpha(X_t - x_0) + \eta_t \tag{13}$$

where $X_t$ is the projection at time bin t, $\alpha$ is the retraction coefficient characterizing the depth of the attractor basin, $x_0$ is the fixed point of the undriven system and $\eta_t$ is zero-mean Gaussian white noise. A larger $\alpha$ reflects a stronger attractor around the fixed point; neural activity retracts more strongly toward $x_o$ per time step. The model was fit by minimizing the negative log-likelihood (NLL):

$$L(\alpha, x_0) = \frac{n}{2}\log(\frac{SSE(\alpha,x_0)}{n}) \tag{14}$$

where SSE is the sum of squared prediction errors across all consecutive bin pairs ($X_t$, $X_{t+1}$) within a session and $n$ is the total number of pairs. The variable $\alpha$ was constrained to a stable retractor regime of (0, 1) by reparametrizing $\alpha = \sigma(\emptyset)$ where $\sigma$denotes the logistic sigmoid. Parameter optimization was initialized at $\emptyset_0 = 0$ (corresponding to $\alpha = 0.5$) with the fixed point initialized at the mean of the observed $X_t$ values. Ten random restarts were used and the solution with the lowest NLL was retained.

For each session, a separate dynamics model was fit for completed trials and abort trials. Consecutive bin pairs ($X_t$, $X_{t+1}$) of neural responses were collected separately for trial categories. Abort trials consisted of non-first aborts only (second abort in a consecutive run onward), excluding transitional first-abort trials and single aborts.

*Existence of distinct attractor basins:* To test whether completed and abort trial ITIs exhibited distinct attractor dynamics, the model was fit separately to completed and aborted trial groups, yielding one $\alpha_C$ and one $\alpha_A$ per session. Sessions were included in analysis if the minimum number of consecutive bin pairs ($X_t$, $X_{t+1}$) in either category exceeded 20. One-sample t-tests against zero were used to determine whether basins were present and a paired t-test was used to determine whether the completed trial basins were deeper than abort basins. The separation of fixed points ($x_{0\_C}, x_{0\_A}$) was assessed with a two-sided paired t-test.

*Shuffle testing:* To confirm that fitted retraction coefficients reflected genuine temporal autocorrelation in the neural dynamics, a within-category shuffle control was performed. For each session and trial category, the vector of $X_t$ values across all consecutive bin pairs was permuted using a derangement procedure such that $X_t$ and $X_{t+1}$ came from different trials of the same condition to destroy the predictive relationship between $X_t$ and $X_{t+1}$ but to retain the marginal distributions of $X_t$ within trial categories. The model was then refit on the shuffled pairs using identical procedures. This was repeated for 25 permutations per session, and the mean shuffled $\alpha$ and $x_0$ across permutations served as the shuffle estimate for that session.

When $X_t$ carries no information about $X_{t+1}$, the model's optimal solution is to ignore $X_t$ and predict the category mean $x_0$ at each step, which results in $\alpha \to 1$ (instantaneous collapse to the fixed point). Real estimates of $\alpha$ were significantly below the shuffle null, and under the shuffle, fitted $\alpha$ converged to 1.0, confirming that the fitted retraction coefficients reflected genuine temporal structure and not an artifact of the pair collection procedure or marginal distributions. The fixed point estimates were not significantly changed by the shuffle and remained near the same values as the real fits for both trial categories. This is expected as $x_0$ is determined by where the category mean lies along the axis discriminating aborts and completed trials and is insensitive to temporal ordering.

*State value modulation of basin depth:* To test whether attractor depth scaled with motivational state value (SV) in a dissociable manner across completed and abort basins, a modified version of the model was used, in which trials within each basin were split into low-SV and high-SV groups sharing a common fixed point but with separate retraction coefficients:

$$X_{t+1} = X_t - \alpha_k(X_t - x_0) + \eta_t, \quad k \in \{low\ SV, high\ SV\} \tag{15}$$

Within each session and basin type, SV outliers were removed (SV≤ 0, Tukey method 1.5 x IQR). Trials in the bottom 100/Npercentile of the inlier SV distribution were assigned to the low-SV group; trials in the top 100/Npercentile were assigned to the high-SV group; middle trials were excluded. N=10 was used to collect 10% splits and capture each end of the ranges of state values. Sessions that had less than 250 pairs (trials and bins) were excluded from analysis. The shared $x_0$ model was fit by minimizing the pooled NLL across both SV groups jointly:

$$L(\alpha_{SVLow}, \alpha_{SVHigh}, x_0) = \frac{n_{SVLow} + n_{SVHigh}}{2} \log(\frac{SSE_{SVLow} + SE_{SVHigh}}{n_{SVLow} + n_{SVHigh}}) \tag{16}$$

This yielded four retraction coefficients per session: $\alpha_{C\_SVLow}, \alpha_{C\_SVHigh}, \alpha_{A\_SVLow}, \alpha_{A\_SVHigh}$ and two shared fixed points $x_{0\_C}$ and $x_{0\_A}$.

Two tests were performed to assess statistical significance of SV on basin depth. First, per-session difference scores were computed for the subset of sessions with both valid basins (n=42): $\Delta\alpha_{C\_SV} = \alpha_{C\_SVHigh} - \alpha_{C\_SVLow}$ and $\Delta\alpha_{A\_SV} = \alpha_{A\_SVHigh} - \alpha_{A\_SVLow}$. A one-sided paired t-test was used to test whether $\Delta\alpha_{C\_SV} > \Delta\alpha_{A\_SV}$. A second test was a linear mixed effects model (LME) fit to all valid session-by-condition observations pooled across both monkeys:

$$\alpha \sim 1 + SV + basin + SV * basin + (1|session) \tag{17}$$

where SV was effect-coded (-0.5 = low, + 0.5=high) and basin was effect coded (-0.5 = abort, +0.5=CT). The interaction term tested whether the effect of SV on $\alpha$ differed between completed and abort basins, specifically whether high SV strengthened the completed trial attractor while weakening the abort attractor. Sessions with only one valid basin contributed only two observations to the model instead of the maximum four; the LME allowed for the inclusion of more sessions with partial data and confirmed the results from the difference score test.

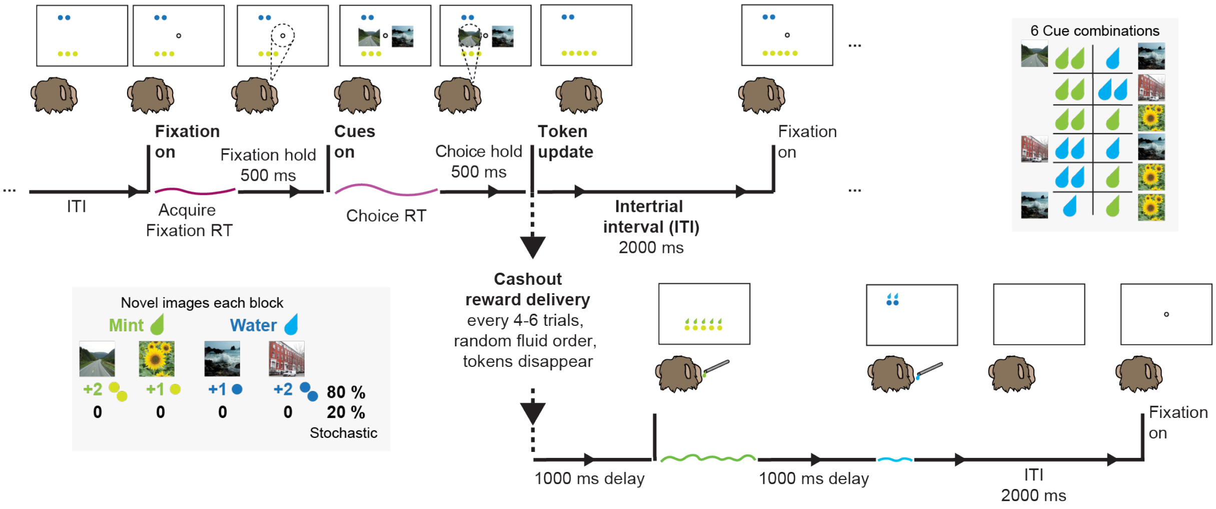


**Fig. S1. Dual fluids tokens task structure and trial timing details.** Two rhesus macaques performed a dual-fluid token bandit task across daily sessions. At the start of each block of 108 trials, four novel images were introduced; each image was associated with a token outcome (+2 Mint, +1 Mint, +1 Water, or +2 Water). The four images defined six cue pair combinations, which were presented pseudorandomly across trials within each block. Each trial, excluding the very first trial of the session, began with a 2000 ms intertrial interval (ITI), during which the accumulated token display remained visible. The monkey then had 2000 ms to acquire a central fixation spot and was required to hold gaze within the fixation window for 500 ms upon acquiring fixation. After a successful fixation hold, two cue images appeared on the screen. The monkey indicated a choice by making a saccade to one image and their holding gaze on the cue for 500 ms. After a successful choice hold, the corresponding number of tokens was added to the screen on 80% of trials; no token change occurred on the remaining 20%. Tokens accumulated across trials and remained visible throughout the session. Cashout occurred randomly every four to six trials. During cashout, one drop of fluid per accumulated token was delivered for each fluid, and each token disappeared as one drop was delivered until all tokens were removed from the screen. Fluid deliveries were separated by a delay of 1000 ms and delivered in a random order. Cashout was followed by the 2000 ms ITI.

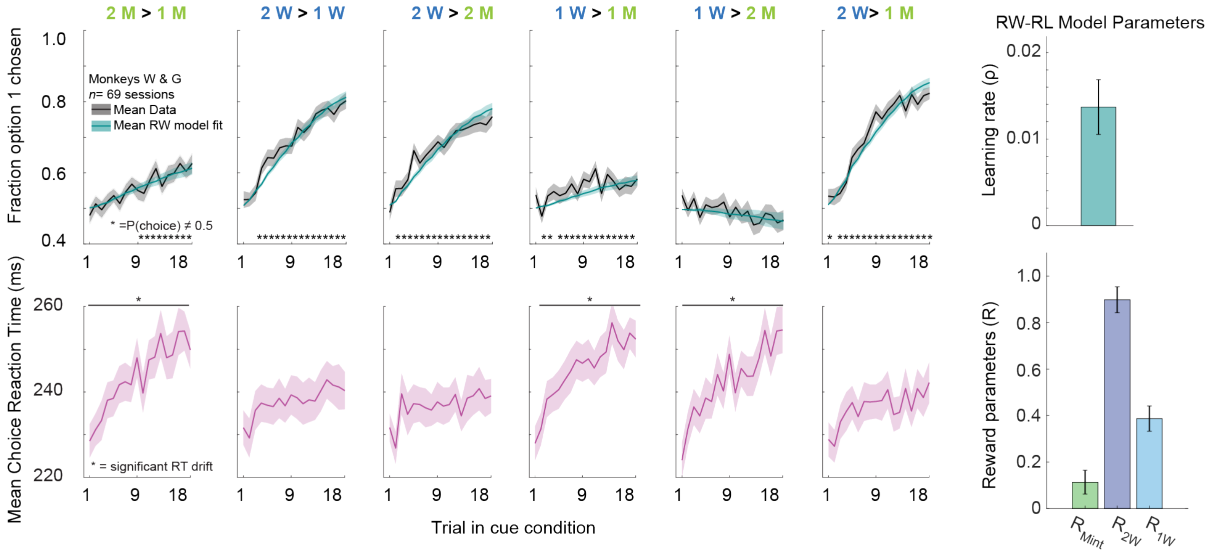


**Fig. S2. Behavioral performance and reaction times across cue conditions.** *Top*: mean fraction of trials on which option 1 of the cue condition was chosen across trials averaged across n=69 sessions from both monkeys (black), overlaid with the average choice probability computed from the Rescorla-Wagner (RW) model fit separately to each session (teal). Asterisks in each panel indicate trials at which choice probability differed significantly from chance (BH-FDR corrected, $p < 0.05$). Error bars: s.e.m. across sessions.

*Bottom row*: mean choice reaction time across trials within each cue condition for the same sessions. Asterisks indicate conditions with a significant drift in reaction times across trials within the block. *Right:* RW model parameters from per-session fits, averaged across n=69 sessions (mean ± SEM across sessions). The learning rate (ρ, top) reflects the rate at which choice probability updated with token outcomes. Fluid-specific reward parameters (R, bottom) reflect the relative subjective value assigned to each fluid type derived from the model fits.

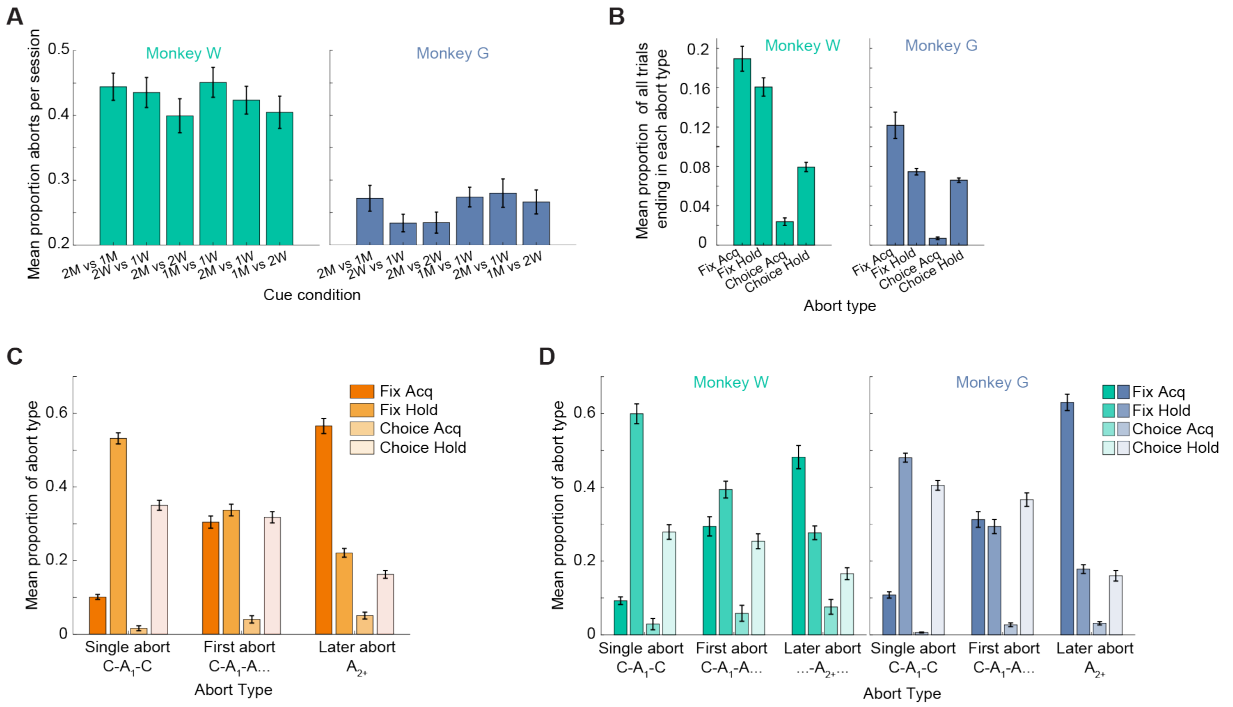


**Fig. S3. Characterization of abort behavior by cue condition, trial epoch, and sequential position across monkeys.**

**(A)** Mean proportion of aborts by cue condition. Abort rate showed a small but significant effect of cue condition in both monkeys (RM-ANOVA, Greenhouse-Geisser corrected: Monkey W: $F(3.64, 105.68) = 3.00$, $p = 0.026$, n=30 sessions; Monkey G $F(4.52, 171.63) = 2.52$, $p = 0.037$, n=39 sessions). No single cue condition dominated abort behavior in either monkey.

**(B)** Mean proportion of aborts by abort type. Fixation aborts were more frequent than choice aborts. FixAcq = failed to acquire fixation, Fix Hold= failed to hold fixation, Choice Acq= failed to acquire a choice image, Choice Hold= failed to hold choice image. Error bars s.e.m.

**(C)** Proportions of aborts by Single abort ($C\text{-}A_1\text{-}C$), First abort ($C\text{-}A_1\text{-}A\ldots$), and Later abort ($A_{2+}$) combined across monkeys. Single aborts were dominated by hold-period errors (Fix Hold + Choice Hold) relative to acquisition errors (Fix Acq + Choice Acq; paired t-test, pooled sessions, $t(68) = 44.70$, $p < 0.0001$). First aborts were distributed across trial epochs except Choice Acq. Across sequential aborts, the proportion of Fix Acq errors increased from First abort to Later abort (paired t-test, $t(68) = -18.20$, $p < 0.0001$, n=69 pooled sessions), consistent with aborts pushing earlier in the trial as disengagement progresses.

**(D)** Same as (C) by monkey. Error bars: s.e.m. Monkey W: 30 sessions, Monkey G: 39 sessions.

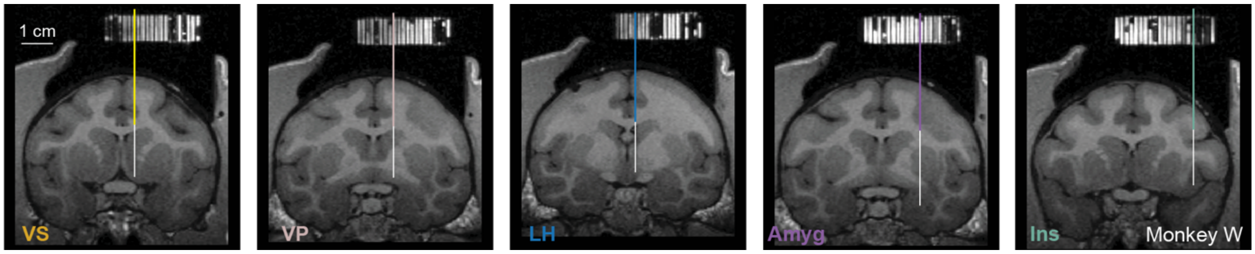


**Fig. S4. Example MRI recording trajectories in Monkey W**. Coronal MRI sections from Monkey W showing electrode targeting for each of the five recording areas. A recording grid was filled with gadolinium contrast agent prior to imaging, producing the bright grid visible at the top of each section. Colored lines indicate the guide tube positions for one set of recordings; white lines indicate example electrode trajectories used to target each area. Sections are shown at the anterior-posterior level appropriate for the approximate center of each structure. Scale bar: 1cm.

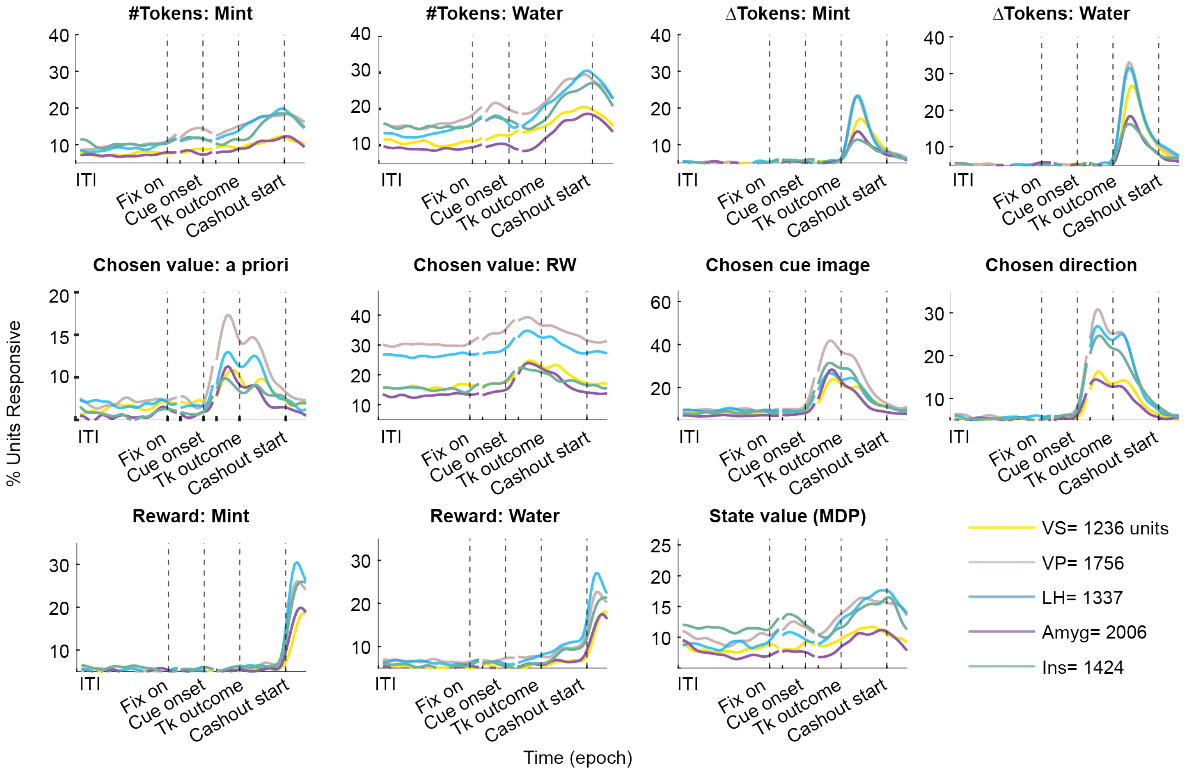


**Fig. S5. Distributed encoding of task features across limbic areas.** Percentage of units with significant modulation by each task variable across trial epochs, combined across monkeys, for each of the five recording areas. Significance was assessed by sliding window multi-way ANOVA models. Dashed vertical lines indicate task epochs. Modulation by all task features was distributed across areas throughout the trial, with no feature restricted to a single limbic structure. RW=Rescorla-Wagner derived chosen value, a priori value= +2 or +1. Data split by individual monkey shown in Supplementary Figure 6.

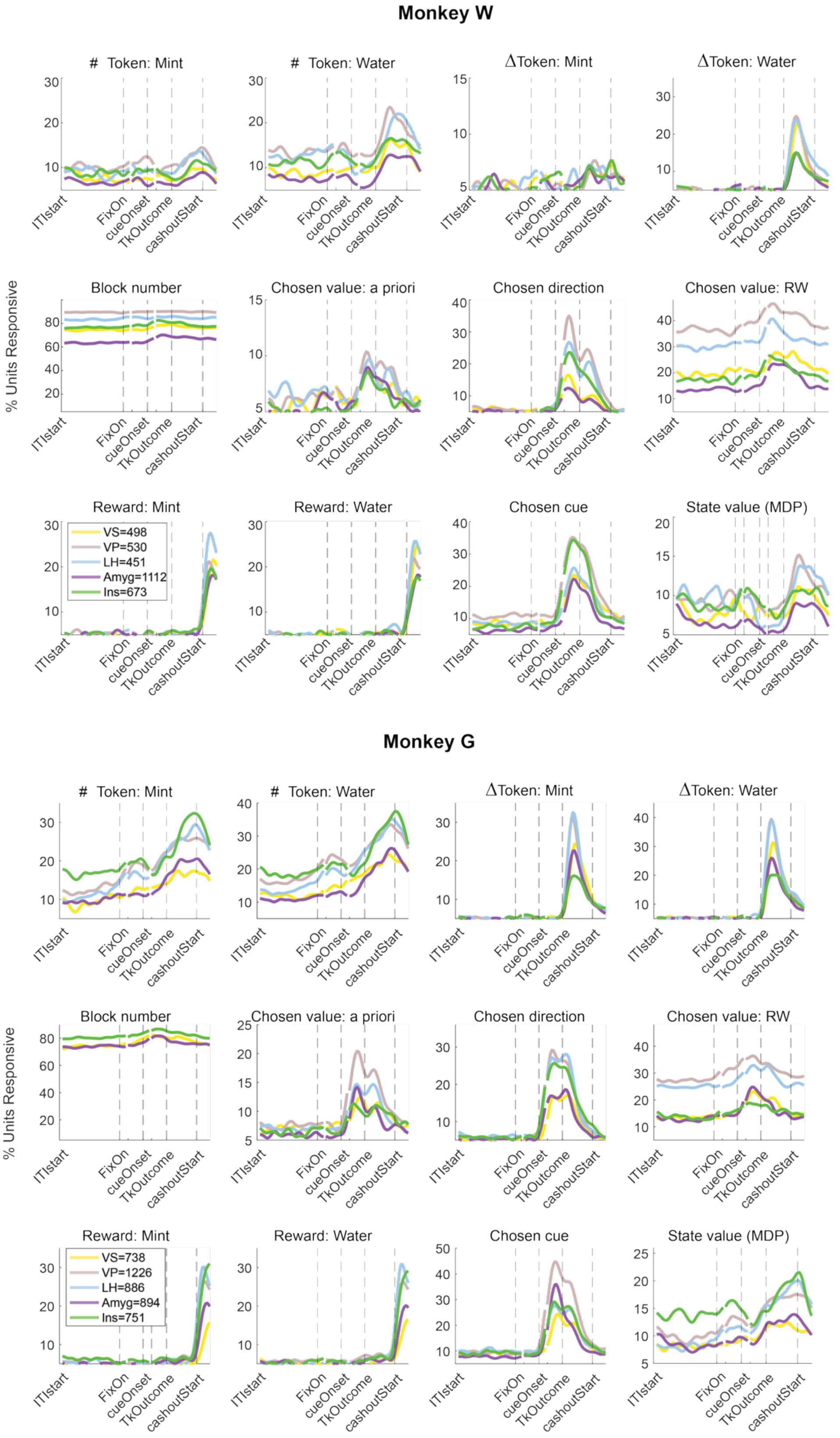


**Fig. S6. Distributed encoding of task features across limbic areas by monkey.** As in Supplementary Figure 5, but shown separately for Monkey W and Monkey G.

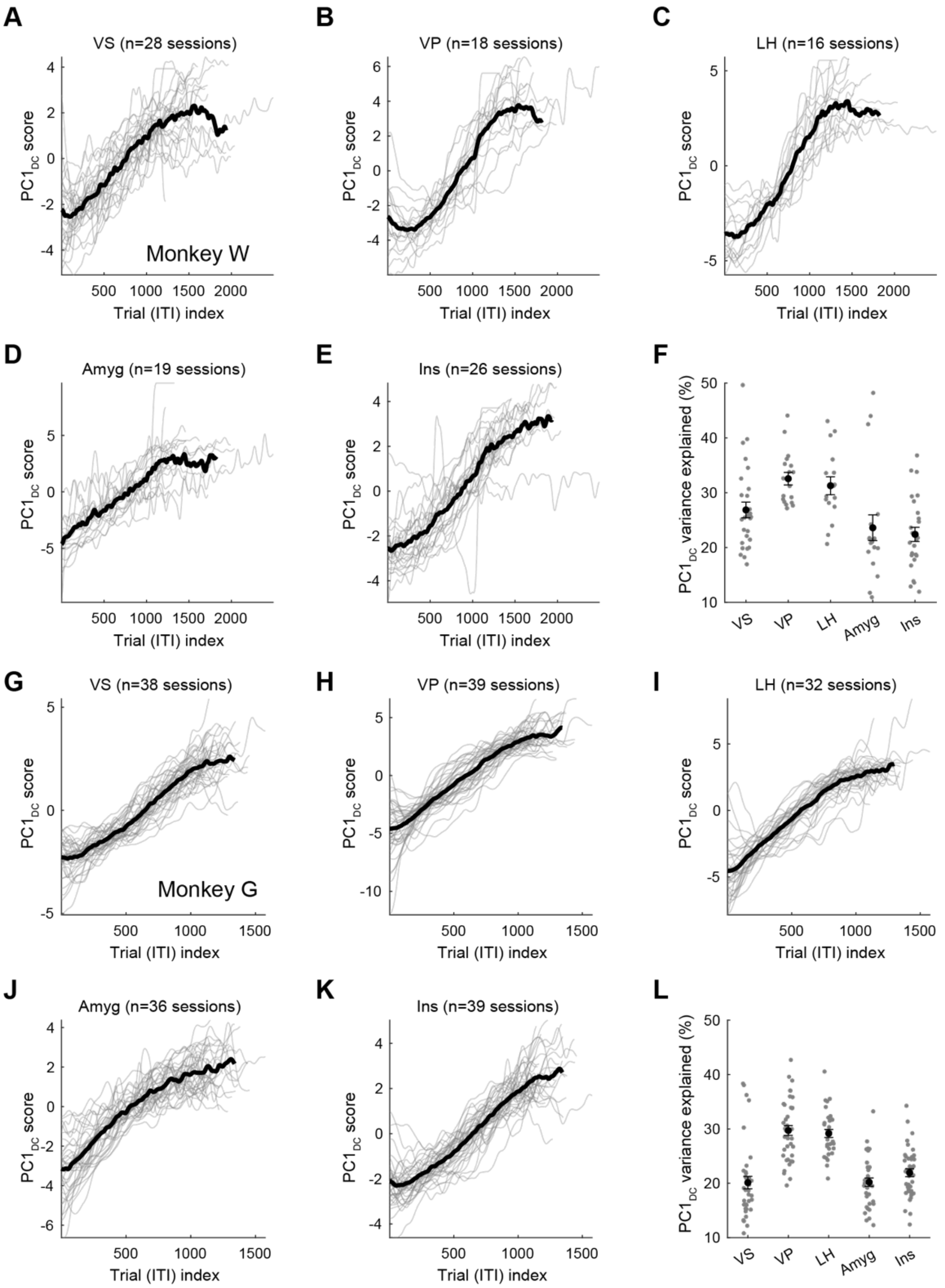


**Fig. S7. Session drift captured by PC1$_{DC}$ of trial-mean ITI population activity.**
**(A-F)** Monkey W. (A-E) PC1$_{DC}$ projection scores across trials for each recording area (gray lines: individual sessions; black line, mean across sessions). PC1 was identified as the component most correlated with trial index and time in session.(DC= Drift Correction)
**(F)** Variance explained by PC1$_{DC}$ per area across sessions (dots: individual sessions; error bars: s.e.m.).
**(G-L)** Same as A-F for Monkey G. PC1$_{DC}$ projections increased monotonically across trials in both monkeys and all areas, consistent with coordinated slow drift. The PC1$_{DC}$ loading was removed from population activity before decoding and subsequent analyses.

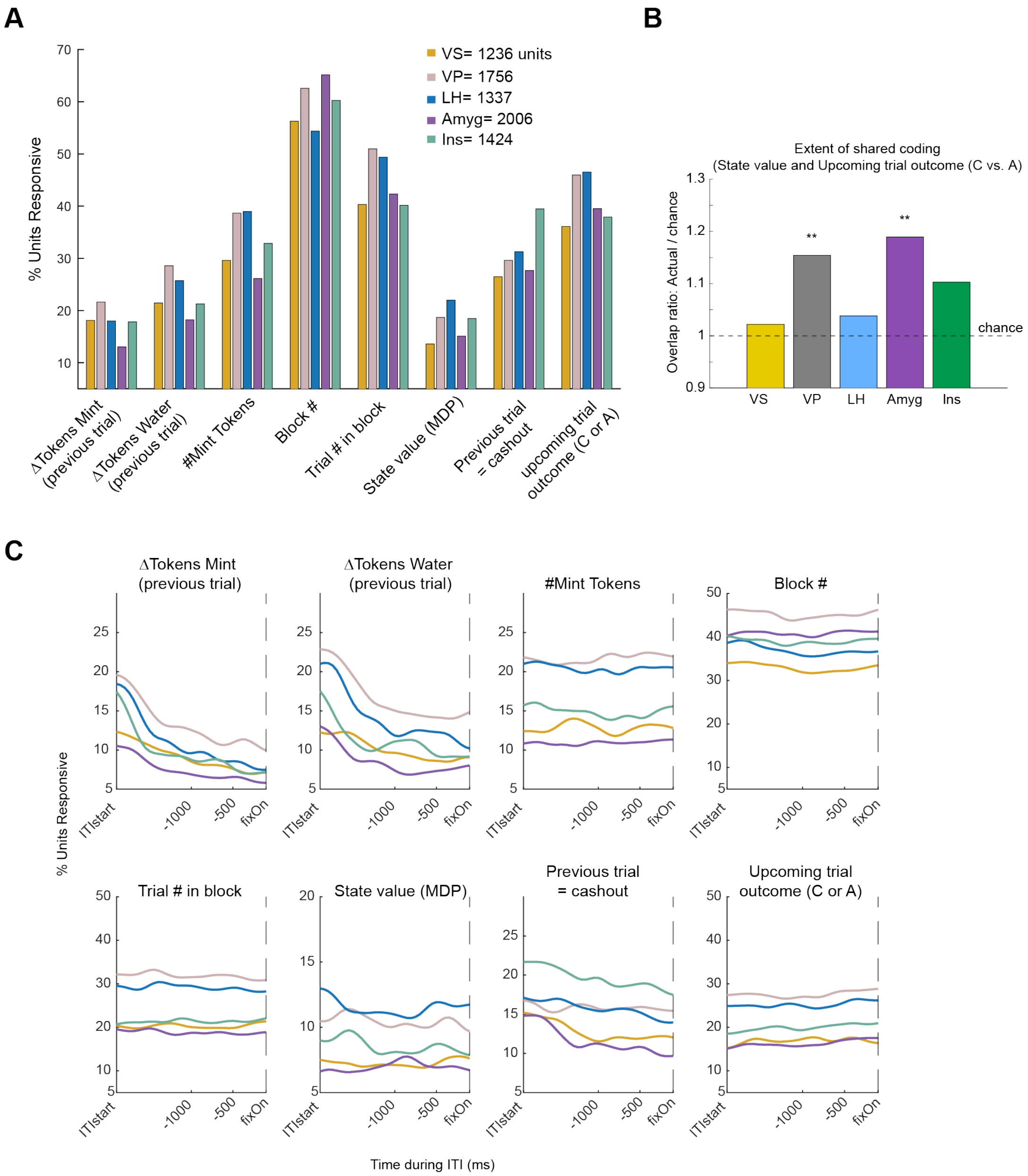


**Fig. S8. Single-unit modulation during the ITI.**

**(A)** Percentage of units per area with significantly different mean ITI firing rates across task features relevant to the ITI, combined across monkeys.

**(B)** Enrichment of single units co-encoding state value and upcoming trial outcome, combined across monkeys. Bars show the ratio of the observed number of co-encoding units to the number expected if the two signals were distributed independently across the population; values above the dashed line (chance) indicate more co-encoding neurons than predicted by chance. Observed versus expected co-encoding units: VS: 62 vs 60.7, p= 0.82; VP: 174 vs 150.7, p= 0.0043; LH: 142 vs 136.8, p = 0.49; Amyg: 142 vs 119.4, p = 0.0039; Ins, 110 vs 99.7, p = 0.15. Co-encoding exceeded chance in VP and Amyg (asterisks indicate significance, chi-square test, Bonferroni-corrected)

**(C)** Same as (A) but computed across the ITI window (200ms bins, 50 ms sliding window).

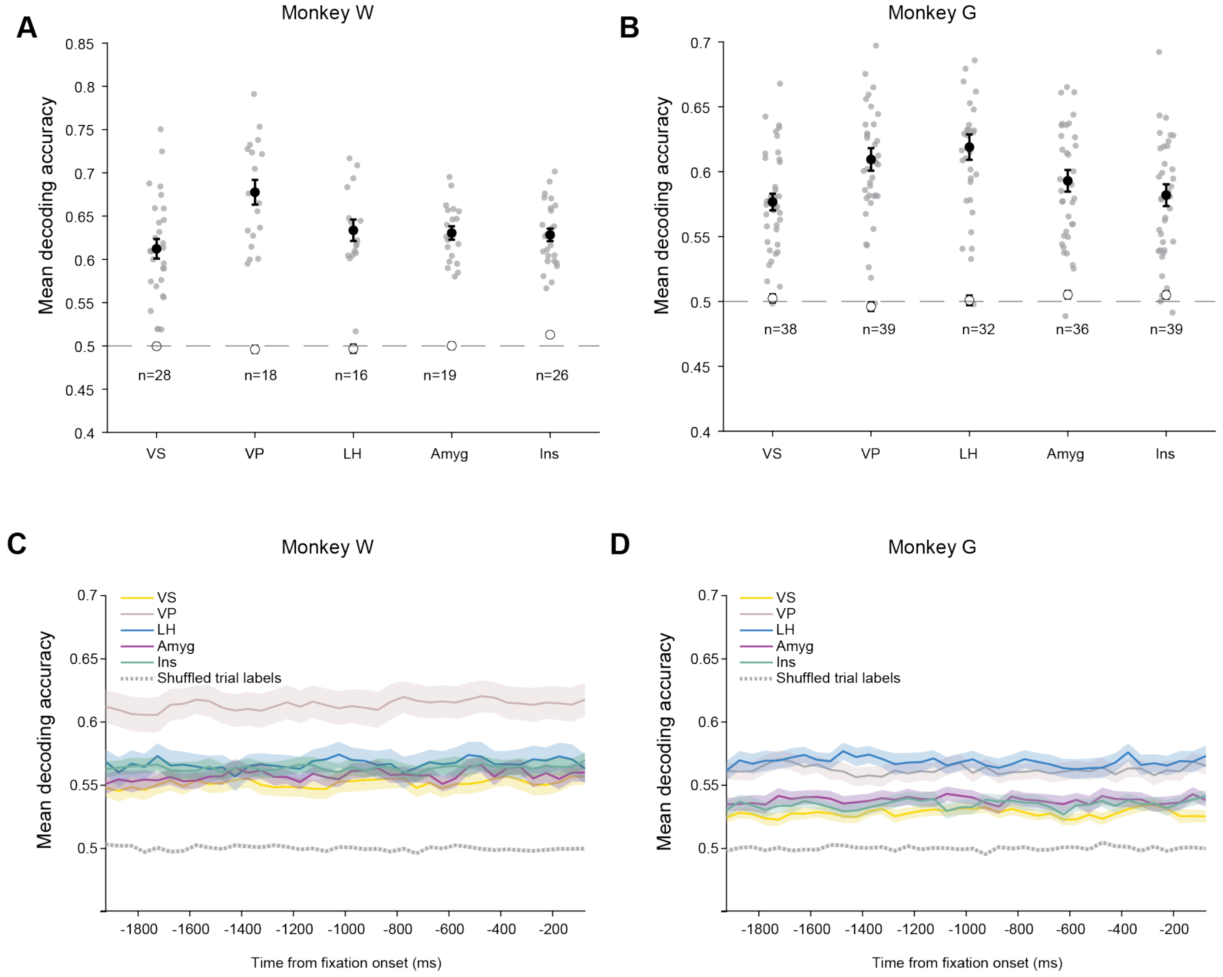


**Fig. S9. ITI decoding accuracy by monkey and across the ITI window.**
**(A, B)** Cross-validated mean decoding accuracy for upcoming abort versus completed trial from mean ITI population activity, shown separately for Monkey W (A) and Monkey G (B). Open circles: shuffled-label chance; dashed line: theoretical chance at 0.5. Session counts per area indicated.
**(C, D)** Decoding accuracy across the ITI time window for Monkey W (C) and monkey G (D), computed in 150 ms bins, sliding window 50 ms, aligned to fixation onset. Colors indicate recording area; dashed line: shuffled-label chance. Decoding performance was stable across the ITI window in both monkeys, with no evidence of an abrupt onset of the motivational signal prior to fixation.

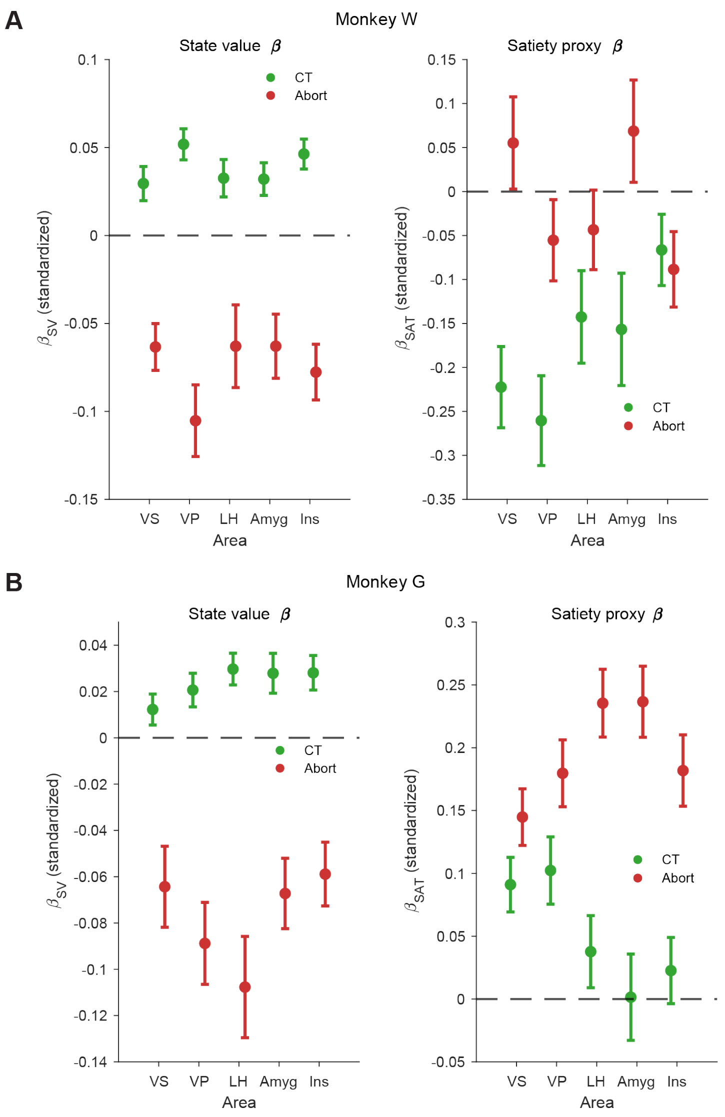


**Fig. S10. Posterior probability regression coefficients for state value and satiety proxy, by monkey.**

**(A, B)** Standardized regression coefficients for state value (left) and satiety proxy (right) from the best fitting full model for Monkey W (A) and Monkey G (B), shown separately for completed trials (green) and abort trials (red). Coefficients reflect the change in posterior probability in standard deviation units per one standard deviation change in each predictor, fit separately per session and area. Error bars: s.e.m. across sessions; full statistics in table S2. Model selection and satiety proxy operationalization are described in the Methods. State value coefficients were significant and in the expected direction across all areas and both trial types, replicating the pooled result in Fig. 3H. Due to drift correction, satiety proxy coefficients were near zero for Monkey W, indicating that session progression did not independently explain the decoded motivational signal. For Monkey G, satiety proxy coefficients were weakly positive for abort trials but near zero for completed trials, suggesting a mild residual correlation between abort-trial posteriors and session progression that likely reflects a genuine motivational signal of satiety accumulation rather than electrode or firing rate drift.

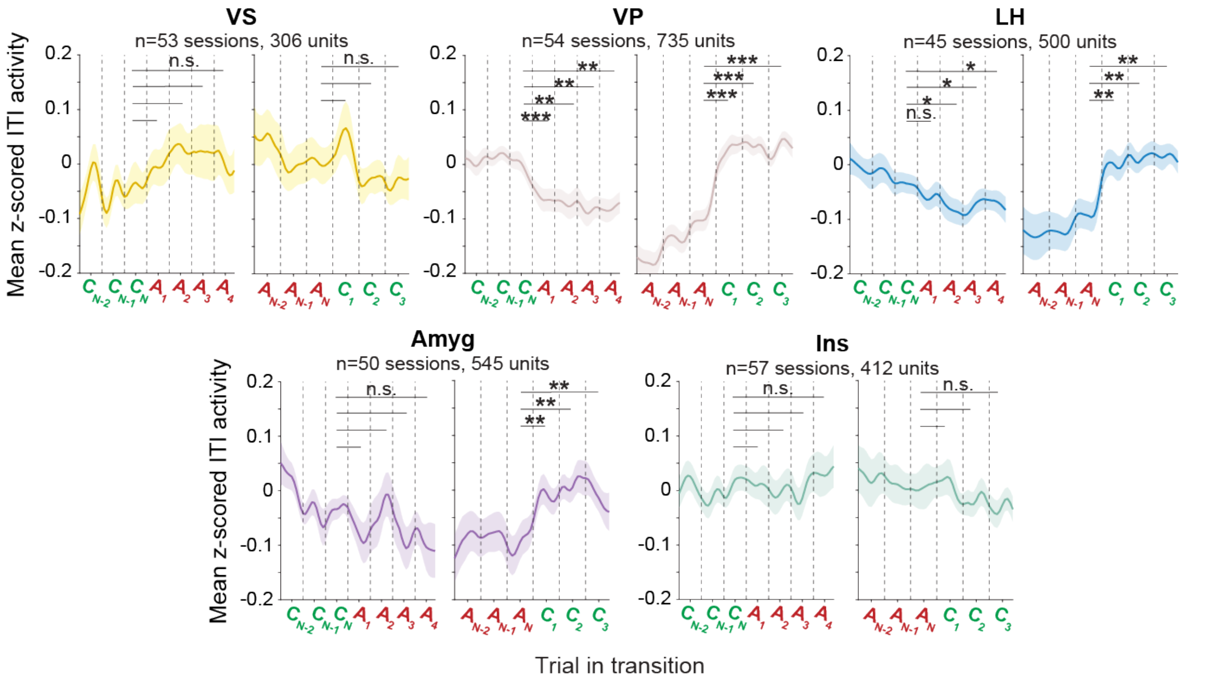


**Fig. S11. Mean ITI firing rates of engagement-modulated units across disengagement and re-engagement transitions.** Mean z-scored ITI activity of units with significant engagement-related responses (abort vs. completed trial, fig. S8), shown across sequential trial transitions spanning the disengagement and re-engagement transitions. Data pooled across monkeys; session and unit counts indicated per panel. Shading: s.e.m. across sessions. Asterisks indicate significant rate shifts between trial positions and level of significance. VP and LH showed significant rate decreases during disengagement, with VP reaching significance at $A_1$ ($t(51)= -3.685$, $p=0.001$) and LH at A2 ($t(44)=-1.972$, $p=0.050$), and significant rate increases at $C_1$ during re-engagement (VP: $t(53)=6.888$, $p<0.0001$; LH: $t(44)=3.310$, $p=0.002$). VS rates increased slowly across the abort run but did not reach significance at any transition, with no significant change at re-engagement. Amygdala showed a significant rate change only during re-engagement ($t(49)=2.574$, $p=0.013$), and Insula showed no significant rate transitions at either transition despite carrying a significant decoding signal in both contexts.

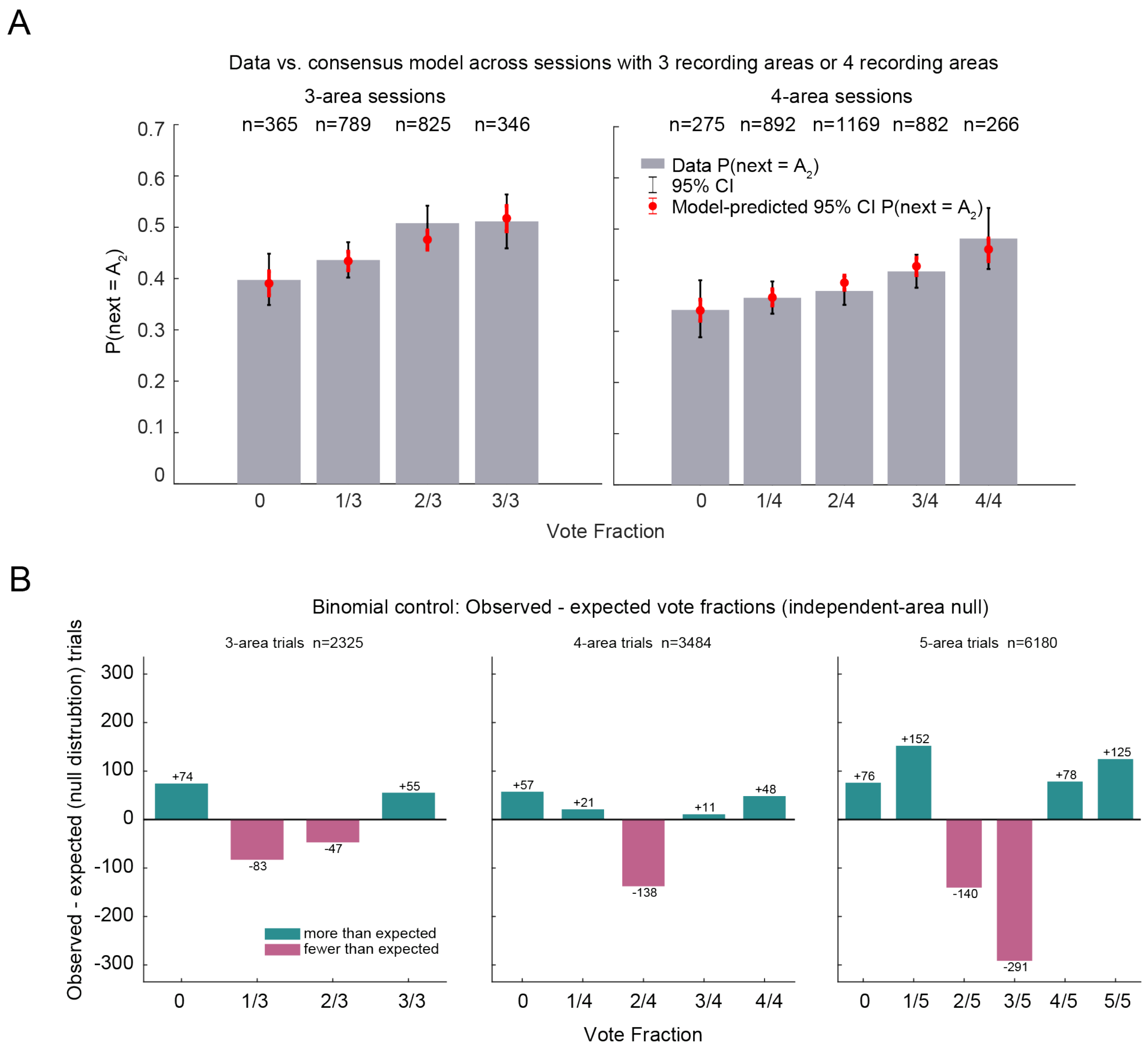


**Fig. S12. Robustness of the consensus results across recording configurations and against an independent-area binomial null.**

**(A)** Consensus model generalization to 3-area (left) and 4-area (right) recording sessions. Bars show observed p(next=$A_2$) per vote fraction bin; red points and error bars show the 95% CI predicted by the GLMM fit simultaneously to all three session types (3-, 4-, and 5-area sessions). Trial counts per bin indicated above bars.

**(B)** Observed minus expected vote-count frequencies under a binomial null assuming independent, equal-probability votes (p=0.5 per area) for 3-area sessions ($\chi^2(3) = 40.0$, p= $1.07*10^{-8}$, n=2325 trials), 4-area sessions ($\chi^2(3) = 40.9$, p= $2.87*10^{-8}$, n=3484 trials), 5-area sessions ($\chi^2(3) = 195.1$, p<0.0001, n=6180 trials). Teal bars indicate vote fractions observed more frequently than the binomial null predicted; pink bars indicate vote fractions observed less frequently. Departures from the binomial null indicate that notes were not statistically independent across areas, consistent with coordinated rather than independent displacement across limbic areas.

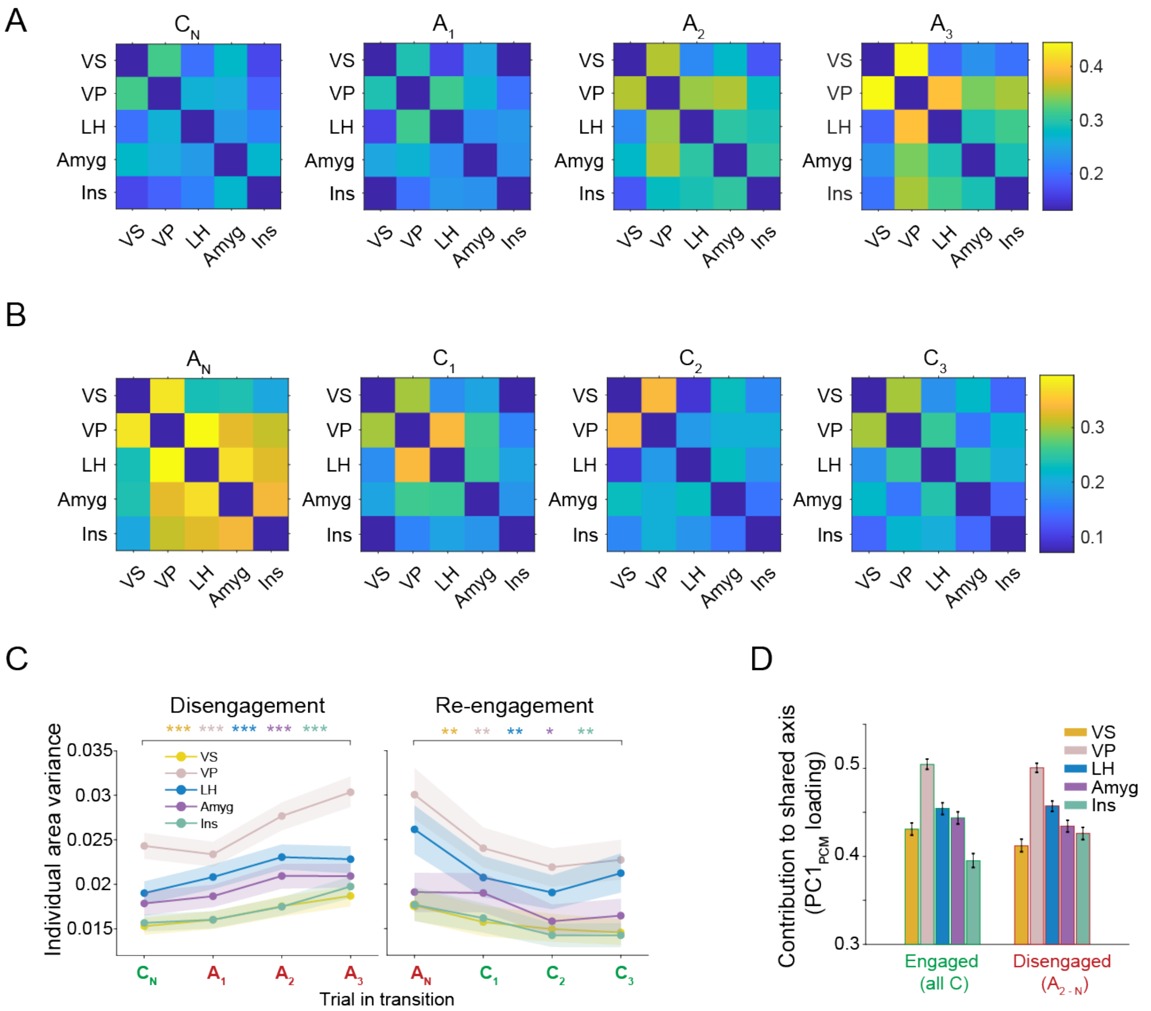


**Fig. S13. Structure and components of inter-area posterior coherence during motivational transitions.**

**(A)** Mean posterior correlation matrices across brain areas at each position in the disengagement transition ($C_N$ through $A_3$, n=2770 pooled sequences of trials) across monkeys and sessions. Color indicates strength of Pearson correlation.

**(B)** Same as (A) for the re-engagement transition ($A_N$ through $C_3$, n=1093 pooled sequences of trials)

**(C)** Individual area posterior variance extracted from the diagonal of the covariance matrix computed across the same trial sequences as (A), (B) and Fig. 5A, increased significantly across the disengagement sequence ($C_N \rightarrow A_3$ $p < 0.0001$ all areas) and decreased significantly during re-engagement ($A_N \rightarrow C_3$: VS: p=0.005, VP: p=0.005, LH: p=0.005, Amyg: p=0.022, Ins: p=0.005). Shaded regions indicate 95% bootstrap confidence intervals.

**(D)** Area loadings onto the first principal component from dimensionality reduction on the posterior correlation matrix ($PC1_{PCM}$) for all engaged trials (n=61324 completed trials) and disengaged trials (n=20246 later abort trials ($A_{2+}$)).

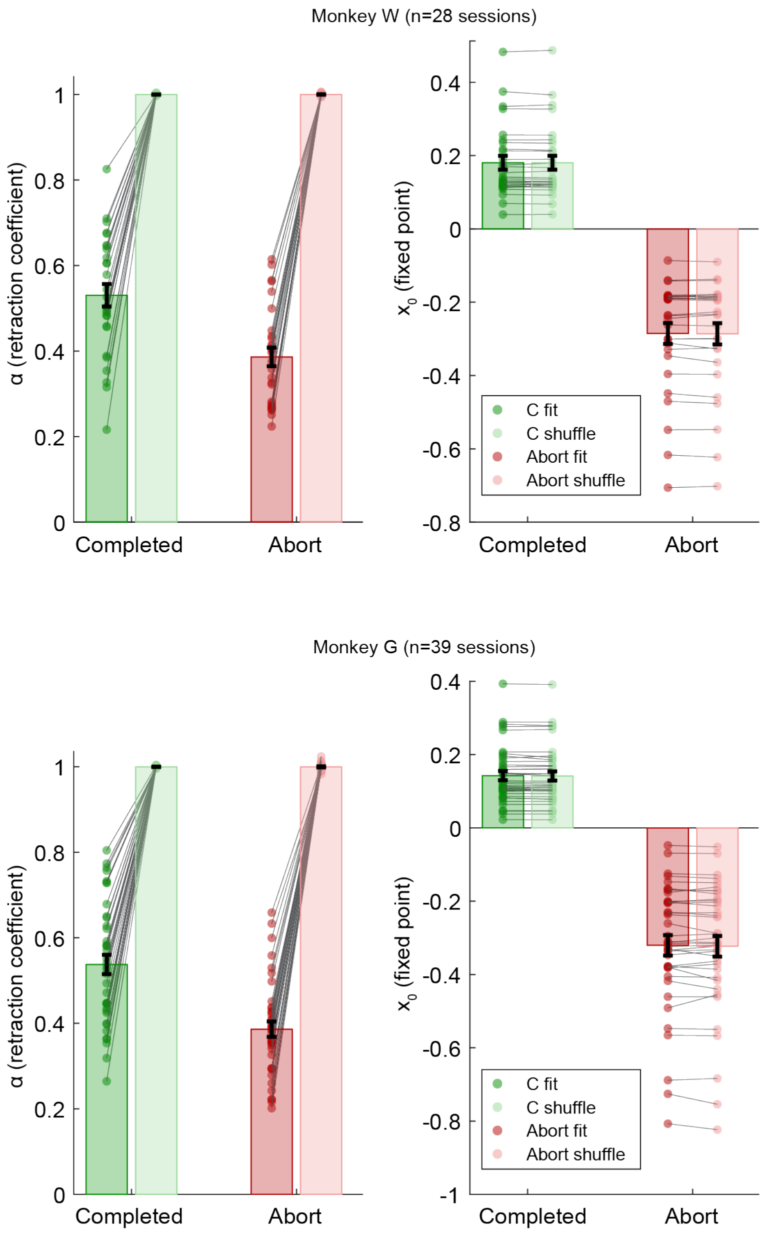


**Fig. S14. Basin fits by monkey and shuffle controls.** Attractor model fits to completed trials and late abort trials for Monkey W (top, n=28 sessions) and Monkey G (bottom, n=39 sessions). Left panels: estimated retraction coefficients for engaged and disengaged basins. Darker bars show fits to real data; lighter bars show fits to time-bin-shuffled data. Shuffle controls are expected to produce an $\alpha \approx 1$, reflecting collapse to the mean in the absence of temporal structure, rather than $\alpha \approx 0$. Both basins had retraction coefficients significantly greater than zero (Monkey W: Completed: $t(27) = 20.039$, $p<0.0001$; Abort: $t(27) = 17.753$, $p<0.0001$; Monkey G: Completed: $t(38) = 23.719$, $p<0.0001$; Abort: $t(38) = 21.518$, $p<0.0001$) and significantly below the shuffle result for both monkeys (Monkey W: Completed: $t(27) = -17.680$; Abort: $t(27) = -27.982$; Monkey G: Completed: $t(38) = -20.406$; Abort: $t(38) = -34.567$, all $p<0.0001$). The engaged basin had a higher retraction coefficient than the disengaged basin (paired t-tests, Monkey W: $t(27) = 5.027$, $p < 0.0001$; Monkey G: $t(38) = 6.612$, $p < 0.0001$). Right panels: estimated fixed points ($x_0$) for engaged and disengaged basins. Fixed point positions were significantly separated for both monkeys (Monkey W: $t(27) = 9.958$, $p < 0.0001$; Monkey G: $t(38) = 11.705$, $p < 0.0001$). Grey lines and dots indicate individual sessions. Error bars: s.e.m.

**Table S1.** Number of neurons recorded in each session and area separated by monkey (Monkey W and Monkey G). Sessions were excluded from individual area analyses if they had fewer than 5 neurons in that area. Total session count = 69 sessions. 39 sessions with all 5 areas; 15 sessions with 4 areas; 11 sessions with 3 areas; 2 sessions with 2 areas; 2 sessions with 1 area.

| Session | Monkey | VS | VP | LH | Amyg | Ins |
|---|---|---|---|---|---|---|
| 21424 | W | NaN | NaN | NaN | 54 | NaN |
| 21624 | W | 20 | NaN | NaN | NaN | NaN |
| 22024 | W | 29 | NaN | NaN | 63 | NaN |
| 22224 | W | 14 | NaN | NaN | 34 | NaN |
| 31524 | W | 17 | 29 | 14 | 45 | 28 |
| 31824 | W | 14 | 27 | 19 | 56 | 21 |
| 32024 | W | 12 | 13 | 15 | 63 | 22 |
| 32124 | W | 24 | 14 | 23 | 80 | 29 |
| 32524 | W | 1 | 20 | 29 | 76 | 31 |
| 32724 | W | 5 | 22 | 21 | 61 | 31 |
| 32924 | W | 18 | 36 | 34 | 59 | 14 |
| 40424 | W | 15 | 23 | 25 | 51 | 5 |
| 52224 | W | 25 | NaN | 37 | 30 | 48 |
| 52424 | W | 7 | NaN | 44 | 46 | 33 |
| 52824 | W | 7 | NaN | 38 | 77 | 41 |
| 53024 | W | 16 | NaN | 33 | 79 | 25 |
| 60424 | W | 11 | NaN | 31 | 63 | 51 |
| 60624 | W | 18 | NaN | 31 | 69 | 32 |
| 61024 | W | 17 | NaN | 30 | 51 | 20 |
| 61224 | W | 6 | NaN | 27 | 55 | 29 |
| 71524 | W | 28 | 41 | NaN | NaN | 39 |
| 71824 | W | 25 | 43 | NaN | NaN | 29 |
| 72224 | W | 31 | 41 | NaN | NaN | 22 |
| 72424 | W | 18 | 34 | NaN | NaN | 21 |
| 72624 | W | 21 | 37 | NaN | NaN | 16 |
| 72924 | W | 30 | 28 | NaN | NaN | 19 |
| 80124 | W | 14 | 27 | NaN | NaN | 25 |
| 80724 | W | 16 | 33 | NaN | NaN | 24 |
| 80924 | W | 17 | 30 | NaN | NaN | 7 |
| 81224 | W | 22 | 32 | NaN | NaN | 11 |
| Totals | W | 498 | 530 | 451 | 1112 | 673 |
| 21325 | G | NaN | 31 | 32 | NaN | 24 |
| 21425 | G | 8 | 32 | 35 | 23 | 29 |
| 21725 | G | 18 | 29 | 40 | 37 | 20 |

| | | | | | | |
|---|---|---|---|---|---|---|
| 21825 | G | 18 | 19 | 20 | 18 | 15 |
| 21925 | G | 11 | 53 | NaN | 16 | 20 |
| 22025 | G | 7 | 48 | NaN | 21 | 15 |
| 22525 | G | 19 | 32 | 30 | 32 | 31 |
| 22625 | G | 18 | 32 | 30 | 24 | 23 |
| 22725 | G | 23 | 34 | 27 | 16 | 18 |
| 22825 | G | 21 | 31 | 22 | 26 | 16 |
| 30325 | G | 18 | 33 | 26 | 31 | 16 |
| 30425 | G | 22 | 28 | 29 | 28 | 7 |
| 30525 | G | 18 | 24 | 32 | 26 | 13 |
| 30625 | G | 23 | 28 | 27 | 28 | 19 |
| 31125 | G | 15 | 29 | 30 | 27 | 18 |
| 31225 | G | 24 | 31 | 32 | 21 | 23 |
| 31325 | G | 23 | 63 | NaN | 30 | 24 |
| 31425 | G | 21 | 24 | 33 | 29 | 24 |
| 31725 | G | 22 | 57 | NaN | 27 | 20 |
| 31825 | G | 24 | 58 | NaN | 31 | 26 |
| 31925 | G | 21 | 57 | NaN | 24 | 13 |
| 32025 | G | 13 | 28 | 31 | 25 | 24 |
| 32525 | G | 21 | 27 | 27 | 34 | 18 |
| 32625 | G | 23 | 28 | 32 | 43 | 21 |
| 32725 | G | 24 | 29 | 29 | 41 | 18 |
| 32825 | G | 23 | 29 | 34 | 32 | 23 |
| 33125 | G | 23 | 33 | 33 | 34 | 17 |
| 40125 | G | 14 | 26 | 27 | 11 | 19 |
| 40225 | G | 17 | 28 | 31 | 22 | 17 |
| 40725 | G | 21 | 32 | 21 | 36 | 18 |
| 40925 | G | 19 | 32 | NaN | 20 | 19 |
| 41525 | G | 24 | 30 | 27 | 22 | 21 |
| 41625 | G | 23 | 25 | 13 | 20 | 20 |
| 41825 | G | 25 | 23 | 28 | 2 | 24 |
| 42225 | G | 22 | 22 | 26 | 12 | 11 |
| 42425 | G | 19 | 14 | 25 | 6 | 15 |
| 42825 | G | 15 | 14 | 26 | 10 | 23 |
| 43025 | G | 19 | 19 | 18 | 6 | 15 |
| 50225 | G | 19 | 14 | 13 | 3 | 14 |
| Totals | G | 738 | 1226 | 886 | 894 | 751 |

**Table S2.** Statistics for regressors: state value (SV) and time-in-session satiety proxy (SAT) from per-session regressions of ITI decoding posteriors on SV and SAT, reported separately for completed (CT) and abort trials (Abt), per area, and per monkey. One-sample t-tests against zero were applied to per-session β coefficients and FDR-corrected across the ten tests within each section (5 areas × 2 trial categories). βSAT results are presented separately per monkey due to inconsistent direction and significance pattern across monkeys.

| Combined Monkeys: beta SV | | | |
|---|---|---|---|
| Area | Trial category | t(df)=t-stat | Corrected p-value |
| VS | CT | t(65)= 3.441 | 1.02e-03 ** |
| VS | Abt | t(65)= -5.572 | 1.30e-06 *** |
| VP | CT | t(56)= 5.104 | 5.15e-06 *** |
| VP | Abt | t(56)= -6.878 | 5.51e-08 *** |
| LH | CT | t(47)=5.346 | 3.69e-06 *** |
| LH | Abt | t(47) = 5.346 | 2.07e-06 *** |
| Amyg | CT | t(47) = -5.559 | 3.42e-05 *** |
| Amyg | Abt | t(54) = 4.550 | 1.35e-06 *** |
| Ins | CT | t(54) = -5.626 | 1.39e-07 *** |
| Ins | Abt | t(64) = 6.224 | 1.11e-07 *** |
| Monkey W: beta SV | | | |
| VS | CT | t(27) = 3.050 | 6.35e-03 ** |
| VS | Abt | t(27) = -4.757 | 1.46e-04 *** |
| VP | CT | t(17) = 5.851 | 9.62e-05 *** |
| VP | Abt | t(17) = -5.153 | 1.59e-04 *** |
| LH | CT | t(15) = 3.049 | 9.02e-03 ** |
| LH | Abt | t(15) = -2.678 | 1.72e-02 * |
| Amyg | CT | t(18) = 3.450 | 4.09e-03 ** |
| Amyg | Abt | t(18) = -3.452 | 4.09e-03 ** |
| Ins | CT | t(25) = 5.448 | 9.62e-05 *** |
| Ins | Abt | t(25) = -4.906 | 1.46e-04 *** |
| Monkey W: beta SAT | | | |
| VS | CT | t(27) = -4.813 | 4.38e-04 ** |
| VS | Abt | t(27) = 1.054 | 0.3348 |
| VP | CT | t(17) = -5.107 | 4.38e-04 ** |
| VP | Abt | t(17) = -1.199 | 0.3158 |
| LH | CT | t(15) = -2.710 | 0.0537 |
| LH | Abt | t(15) = -0.962 | 0.3515 |
| Amyg | CT | t(18) = -2.455 | 0.0612 |
| Amyg | Abt | t(18) = 1.182 | 0.3158 |
| Ins | CT | t(25) = -1.637 | 0.1904 |
| Ins | Abt | t(25) = -2.065 | 0.0988 |
| Monkey G: beta SV | | | |
| VS | CT | t(37) = 1.826 | 0.0759 |
| VS | Abt | t(37) = -3.677 | 1.07e-03 ** |
| VP | CT | t(38) = 2.846 | 7.89e-03 ** |

| | | | |
|---|---|---|---|
| VP | Abt | t(38) = -5.012 | 1.28e-04 *** |
| LH | CT | t(31) = 4.328 | 2.91e-04 *** |
| LH | Abt | t(31) = -4.924 | 1.33e-04 *** |
| Amyg | CT | t(35) = 3.235 | 3.32e-03 ** |
| Amyg | Abt | t(35) = -4.418 | 2.91e-04 *** |
| Ins | CT | t(38) = 3.770 | 9.25e-04 *** |
| Ins | Abt | t(38) = -4.272 | 2.91e-04 *** |
| Monkey G: beta SAT | | | |
| VS | CT | t(37) = 4.193 | 0.0003 ** |
| VS | Abt | t(37) = 6.431 | 3.29e-07 *** |
| VP | CT | t(38) = 3.818 | 6.90e-04 ** |
| VP | Abt | t(38) = 6.740 | 1.84e-07 *** |
| LH | CT | t(31) = 1.315 | 0.2478 |
| LH | Abt | t(31) = 8.716 | 3.83e-09 *** |
| Amyg | CT | t(35) = 0.042 | 0.9664 |
| Amyg | Abt | t(35) = 8.376 | 3.83e-09 *** |
| Ins | CT | t(38) = 0.859 | 0.4395 |
| Ins | Abt | t(38) = 6.396 | 3.29e-07 *** |

**Table S3. Statistics for slope tests on SVM decoder posteriors across disengagement and re-engagement transitions, reported separately for each monkey.** For each brain area, t(df) denotes the t-statistic from a one-tailed one-sample t-test against zero, with Holm-Bonferroni corrected p-values. One-way ANOVA results testing for differences across areas are reported below each slope type.

| Monkey W: Disengagement Slopes | | | |
|---|---|---|---|
| Area | Slope | t(df)=t-stat | Corrected p-value |
| VS | CT | t(28)= -0.183 | 0.428 |
| VS | Abt | t(28)= 4.076 | 0.0007 *** |
| VP | CT | t(17)= -0.562 | 0.2906 |
| VP | Abt | t(17)= 4.341 | 0.0007 *** |
| LH | CT | t(15)= -0.644 | 0.2646 |
| LH | Abt | t(15)= 2.935 | 0.0051 ** |
| Amyg | CT | t(18)= -1.467 | 0.0799 |
| Amyg | Abt | t(18)= 3.988 | 0.0009 *** |
| Ins | CT | t(25)= -0.548 | 0.2943 |
| Ins | Abt | t(25)= 5.679 | 1.63e-05 *** |
| One way ANOVA across areas (Abort slope): F(4,102)= 2.737, p=0.0328 | | | |
| Monkey G: Disengagement Slopes | | | |
| VS | CT | t(34)= 1.641 | 0.9450 |
| VS | Abt | t(34)= 4.268 | 0.0003 *** |
| VP | CT | t(35)= 0.589 | 0.7203 |
| VP | Abt | t(35)= 3.816 | 0.0008 *** |
| LH | CT | t(29)= 0.179 | 0.5705 |
| LH | Abt | t(29)= 5.337 | 2.48e-05 *** |

| | | | |
|---|---|---|---|
| Amyg | CT | t(32)= -0.494 | 0.3123 |
| Amyg | Abt | t(32)= 3.068 | 0.0035 ** |
| Ins | CT | t(35)= 0.684 | 0.7508 |
| Ins | Abt | t(35)= 3.134 | 0.0035 ** |
| One way ANOVA across areas (Abort slope): F(4,165)= 2.359, p=0.0556 | | | |
| Monkey W: Re-engagement Slopes | | | |
| VS | Abt | t(27)= -3.712 | 0.0009 *** |
| VS | CT | t(27)= 1.496 | 0.2350 |
| VP | Abt | t(17)= -4.992 | 0.0002 *** |
| VP | CT | t(17)= 3.265 | 0.0114 * |
| LH | Abt | t(15)= -3.469 | 0.0017 ** |
| LH | CT | t(15)= 1.611 | 0.2350 |
| Amyg | Abt | t(18)= -4.466 | 0.0004 *** |
| Amyg | CT | t(18)= 1.644 | 0.2350 |
| Ins | Abt | t(25)= -4.767 | 0.0002 *** |
| Ins | CT | t(25)= 1.379 | 0.2350 |
| One way ANOVA across areas (CT slope): F(4,102)= 1.195, p=0.3178<br>One way ANOVA across areas (Abt slope): F(4,102)= 1.480, p=0.2136 | | | |
| Monkey G: Re-Engagement Slopes | | | |
| VS | Abt | t(37)= -2.415 | 0.0104 * |
| VS | CT | t(37)= 2.004 | 0.0391 * |
| VP | Abt | t(38)= -4.349 | 0.0002 *** |
| VP | CT | t(38)= 3.894 | 0.0010 ** |
| LH | Abt | t(31)= -3.803 | 0.0009 *** |
| LH | CT | t(31)= 2.908 | 0.0133 * |
| Amyg | Abt | t(35)= -3.699 | 0.0009 *** |
| Amyg | CT | t(35)= 2.502 | 0.0258 * |
| Ins | Abt | t(38)= -4.521 | 0.0001 *** |
| Ins | CT | t(38)= 2.136 | 0.0391 * |
| One way ANOVA across areas (CT slope): F(4,179)= 0.679, p=0.6072<br>One way ANOVA across areas (Abt slope): F(4,179)= 1.480, p=0.1022 | | | |

**References for Supplementary Materials** (101-107)
